# Key informant perceptions on wildlife hunting in India during the COVID-19 lockdown

**DOI:** 10.1101/2021.05.16.444344

**Authors:** Uttara Mendiratta, Munib Khanyari, Nandini Velho, Kulbhushansingh Ramesh Suryawanshi, Nirmal Kulkarni

**Affiliations:** Wildlife Conservation Society – India, 551, 7th Main Road Rajiv Gandhi Nagar, 2nd Phase, Kodigehalli, Bengaluru - 560 097, Karnataka, India; Nature Conservation Foundation, 1311, “Amritha”, 12th Main, Vijaynagar 1st Stage, Mysore, 570 017 India; Interdisciplinary Center for Conservation Sciences, Oxford University, 11a Mansfield Road, Oxford, OX1 3SZ; Department of Biological Sciences, University of Bristol, 24 Tyndall Avenue, Bristol, BS8 1 TQ; Srishti Manipal Institute of Art, Design and Technology, Yelahanka, Bengaluru, India; Snow Leopard Trust, 4649 Sunnyside Ave N, Suite 325, Seattle, WA 98103, USA; Mhadei Research Centre, 6, Hiru Naik Bldg Dhuler, Mapusa Goa 40507, India

**Author notes:** corresponding author: Email –.

**Keywords:** Lockdown, COVID-19, hunting, food security, bushmeat

## Abstract

Lockdowns intended to control the COVID-19 pandemic resulted in major socioeconomic upheavals across the world. While there were numerous reports of these lockdowns benefiting wildlife by reducing human movement and habitat disturbance, increased hunting during these lockdowns emerged as a conservation concern, particular in tropical Asia and Africa. We used online interviews with key informants including wildlife researchers, enforcement staff and NGO employees (N=99), and media reports (N=98), to examine the impacts of India’s COVID-19 lockdown (March-May 2020) on wildlife hunting across the country. We asked whether and how hunting patterns changed during the lockdown, and explored socioeconomic and institutional factors underlying these changes. Over half the interviewees spread over 43 administrative districts perceived hunting (mammals, in particular) to have increased during the lockdown relative to a pre-lockdown reference period. Interviewees identified household consumption (53% of respondents) and sport and recreation (34%) as main motivations for hunting during the lockdown, and logistical challenges for enforcement (36%), disruption of food supply (32%), and need for recreational opportunities (32%) as key factors associated with hunting during this period. These insights were corroborated by statements by experts extracted from media articles. Collectively, our findings suggest that the COVID-19 lockdown potentially increased hunting across much of India, and emphasize the role of livelihood and food security in mitigating threats to wildlife during such periods of acute socioeconomic perturbation.

## 1.0 Introduction

The COVID-19 pandemic has posed unprecedented challenges to humanity. Starting March 2020, countries across the world attempted to contain transmission of the COVID-19 virus by imposing nationwide lockdowns (Karnon 2020). These lockdowns led to unemployment, income loss, food supply chain disruptions, and impacted people’s daily lives and mental health in myriad of ways (Kochhar et al. 2020; Krause et al. 2020). In India, for example, strict lockdowns during March-May 2020 were associated with widespread unemployment and supply chain disruptions leading to food insecurity – a survey of Indian wage workers found that 80% households consumed less food during the lockdown than before (Kesar et al. 2020). Death and suffering were compounded by the large-scale migrations of urban work forces who embarked on long and arduous journeys to return to their rural homes (Bhattamishra 2020; Srivastava 2020).

Globally, the COVID-19 lockdowns had a number of other impacts including that on wildlife across the globe. On one hand, preliminary reports showed wildlife benefiting from reduced human mobility and habitat disturbance during the “Anthropause” (Rutz et al. 2020). On the other hand, the intensification of natural resource extraction including wildlife hunting during this period (Diffenbaugh et al. 2020), particularly across African and Asian nations was reported (Aditya et al. 2021; Badola 2020; Ghosal and Casey 2020; Manenti et al. 2020). For example, illegal hunting and trade of the pangolins in India (Aditya et al. 2021), and that of the critically endangered Giant Ibis in Cambodia, reportedly spiked during the lockdown (Alberts 2020). In India, where hunting of all wildlife barring a handful of “vermin species” (e.g. certain rodents and bats) is prohibited by law (Wild Life Protection Act 1972), reports of hunting in the media doubled during the lockdown (Badola 2020).

Impacts of pandemics on human societies and the economy are in many ways akin to those of war (Banerjee and Duflo 2020; Gaynor et al. 2020). It might therefore be expected that pandemic-related lockdowns and resultant disruptions of food supply chains might increase the demand for wild meat in landscapes where wildlife is available (Borgerson et al. 2019; Bowlin 2020; Jambiya, Milledge and Mtango, 2007). As in the case of war, the pandemic and lockdown could also hamper the functioning of enforcement agencies responsible for wildlife protection (Troumbis and Zevgolis 2020). For example, if patrolling by field staff is constrained by the lockdown (Humphrey 2020), as it often is by war and civil strife (Dutta 2020; Gaynor et al. 2016), this too could contribute to increased hunting. Thus, documenting the impacts of the COVID-19 lockdown on wildlife hunting and examining the socio-economic and institutional factors that potentially underlie these impacts can help conservation practitioners prepare better for future pandemics, lockdowns, and other such socio-economic shocks.

In this study, we use online surveys of key informants, combined with analyses of news media reports, to explore perception of the COVID-19 lockdown on hunting in India. Given that logistical constraints precluded primary data collection on hunting or interviews with hunters, we interviewed wildlife experts and conservation practitioners who were either stationed within focal landscapes themselves, or were in touch with colleagues and teams stationed in these landscapes, during the lockdown. Specifically, we examined perceptions regarding the impact of the lockdown on: (1) locations, targeted species, and groups responsible for hunting; (2) motivations and other socio-economic factors associated with hunting; and (3) functioning of wildlife law enforcement and other counter-hunting strategies.

## 2.0 Materials and Methods

### 2.1 COVID-19 lockdown in India

The Government of India implemented a strict nationwide lockdown from 24^th^ March to 3^rd^ May 2020, which comprised a first phase from 24^th^ March to 14^th^ April and a second phase from 15^th^ April to 3^rd^ May. This lockdown featured strict regulations that suspended all non-essential economic activity and public transport systems, which greatly reduced movement of people. The cessation of economic activity led to the loss or suspension of employment for millions of migrant workers in urban centres, many of whom travelled thousands of kilometres on foot or by bicycle to return to their rural homes. The strict lockdown was followed by a series of ‘unlocking’ steps over which regulations on economic activity and human movement were lifted in a phased manner.

### 2.2 Questionnaire

An online questionnaire (via Google forms; Supplementary Material 1) was used to record the perceptions of wildlife researchers and conservation practitioners on the impacts of the COVID-19 lockdown (25^th^ March to 3^rd^ June 2020) on wildlife hunting in their respective regions of familiarity within India. The survey was circulated through emails to individuals, institutions and groups associated with wildlife research and conservation, and a snow-ball approach helped expand the key informant network. The survey comprised 12 structured and five open-ended questions on how the lockdown affected (1) patterns of hunting; (2) motivations and factors associated with hunting; and (3) counter-hunting strategies including enforcement (Table 1; Supplementary Material 1). Respondents were only permitted to report for locations at which they were stationed during the lockdown themselves (Direct), or at which colleagues, assistants or collaborators with whom they were in contact were stationed during the lockdown (Indirect; see Question 5 in Supplementary Material 1). The two month prior to the lockdown (23^rd^ January to 24^th^ March 2020) were used as comparison.

**Table 1:** Major objectives of the study along with the corresponding topics covered by questionnaire. (Please refer to Supplementary Material 1 for the complete survey form.)

| Objective | Q. No. | Topics covered |
| --- | --- | --- |
| 1. Patterns of hunting | 4-9, 11 | Locations, target taxa, hunting groups, changes in hunting during lockdown |
| 2. Motivations and factors | 10, 12 | Change in motivations to hunt, factors affecting hunting during lockdown |
| 3. Counter-hunting strategies | 13-16 | Lockdown impacts on enforcement and other counter-hunting strategies |

This survey was reviewed and approved by a research ethics committee at Nature Conservation Foundation (NCF-EC-29/04/2020-(49)) prior to circulation. No personal identification information was included in the survey (Supplementary Material 1) and all data has been anonymised.

A total of 99 key informants responded to the survey (79 male; 20 female), including 64 respondents aged 18-34 and 29 respondents aged 35-54. Key informants identified themselves as working with conservation NGOs (N=45), universities (N=23), government staff (N=12), journalists and researchers (N=10) and commercial enterprises associated with wildlife landscapes such as tourism, agriculture, plantations (n=9). Sixty-four respondents were at the location that they were reporting for, and the information was based on their observations alone (21), or combined with information from colleagues, assistants and collaborators (43). Thirty-five respondents based their responses on information provided to them by colleagues, assistants, and collaborators at location during lockdown. Forty-one respondents had direct sighting or first-hand knowledge of hunting events. Illegal fishing (N=29), presence of snares and traps (N=22), and enforcement action (N=17) were some other indicators of hunting.

### 2.3 Media reports

Using online search-engines, we compiled 98 media articles that reported on hunting during the lockdown from across India. Articles dated between 3 May to 31 May (phase 3 and 4 of the lockdown) were also included given the expected lag in reporting. Search phrases included ‘India’, ‘lockdown’, ‘COVID-19’, ‘wildlife hunting’ and ‘wildlife poaching’. From each article, we extracted and coded statements by experts as responses to questions 8, 10, 12 and 13 of the online survey (Supplementary Material 1). In cases where expert statements could not objectively be assigned to survey question categories, these were coded “Don’t know”. To avoid duplication, we discarded statements by individual experts that were repeated across multiple media outlets – a total of 95 unique statements by 75 experts were thus retained.

### 2.4 Analysis

For the key informant interviews and the coded expert statements from media reports, we calculated the percentage of respondents that selected each response category. For the interviews, we also bootstrapped with replacement (10,000 iterations) and estimated means and 95% confidence intervals. We used a chi-square to explore associations between motivations for hunting and focal taxa, and motivations and lockdown-related factors (see questions 8, 10 and 12 in Supplementary Material 1). We used R 4.0.3 (RStudioTeam 2020) and QGIS 3.6 for our analyse (QGIS 2020).

## 3.0 Results

### 3.1 Questionnaire survey: Patterns of hunting

The 99 unique key-informant responses came from 74 districts across 23 Indian states (Figure 1). Over half of the respondents (56%; 95% CI: 40% – 74%) perceived hunting to have increased during the lockdown relative to the pre-lockdown period, 10% (95 % CI: 5%-16%) reported no change, and 6% (95% CI: 1%-13%) reported a decrease, while 27% (95% CI: 19 – 36) were uncertain (‘Don’t know’) (Supplementary Material 2 Table1). Increased hunting during the lockdown was reported from 43 districts across 19 states, while 15 districts across 11 states either reported no change or a decrease in hunting (Fig. 1).

**Figure 1.**
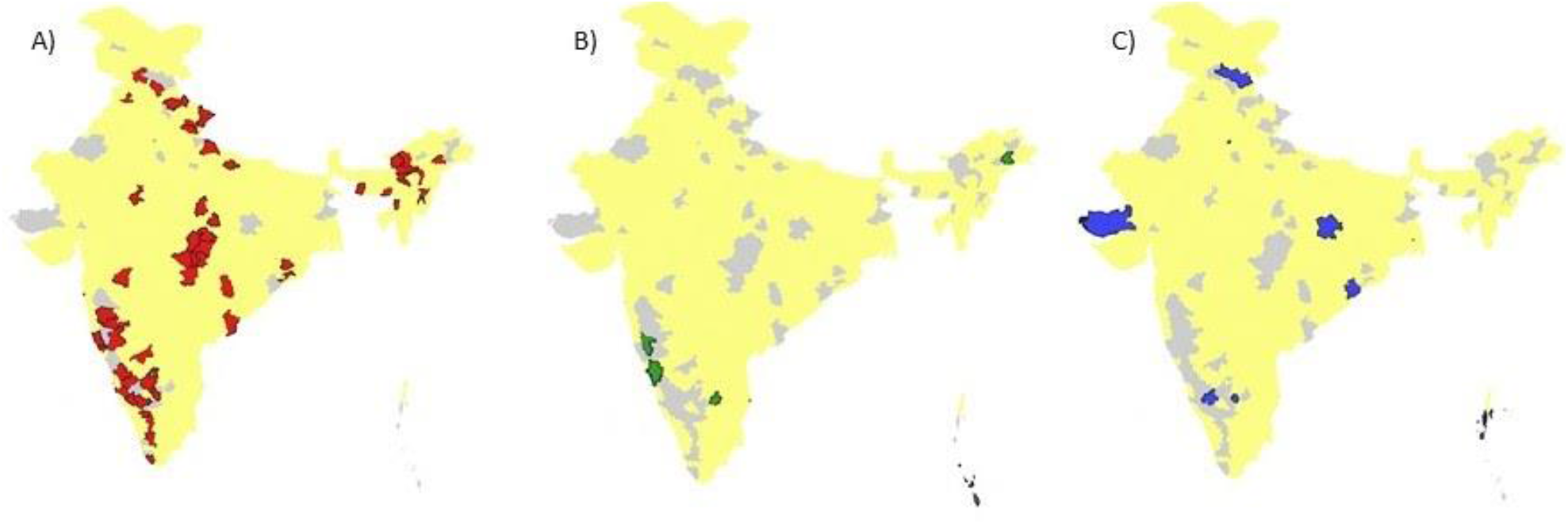
Districts from which data was contributed are marked in grey A) an increase in reports of hunting during lockdown are marked in red (N=43), B) reports of decrease in hunting during lockdown are marked in green (N=6), and C) reports of no change in hunting levels are marked in blue (N=9).

According to the key informants, hunting of mammals (55%; 95% CI: 45% – 66%), fish and crustaceans (43%; 95% CI: 34% – 54%) and birds (35%; 95% CI: 26% – 44%) were higher during the lockdown (Fig.2; Supplementary Material 2 Table 2). For reptiles and amphibians, information on hunting level was sparse, with 34% (95% CI: 25% - 43%) picking “Don’t know” regarding changes in hunting levels (Supplementary Material 2 Table 2).

**Figure 2.**
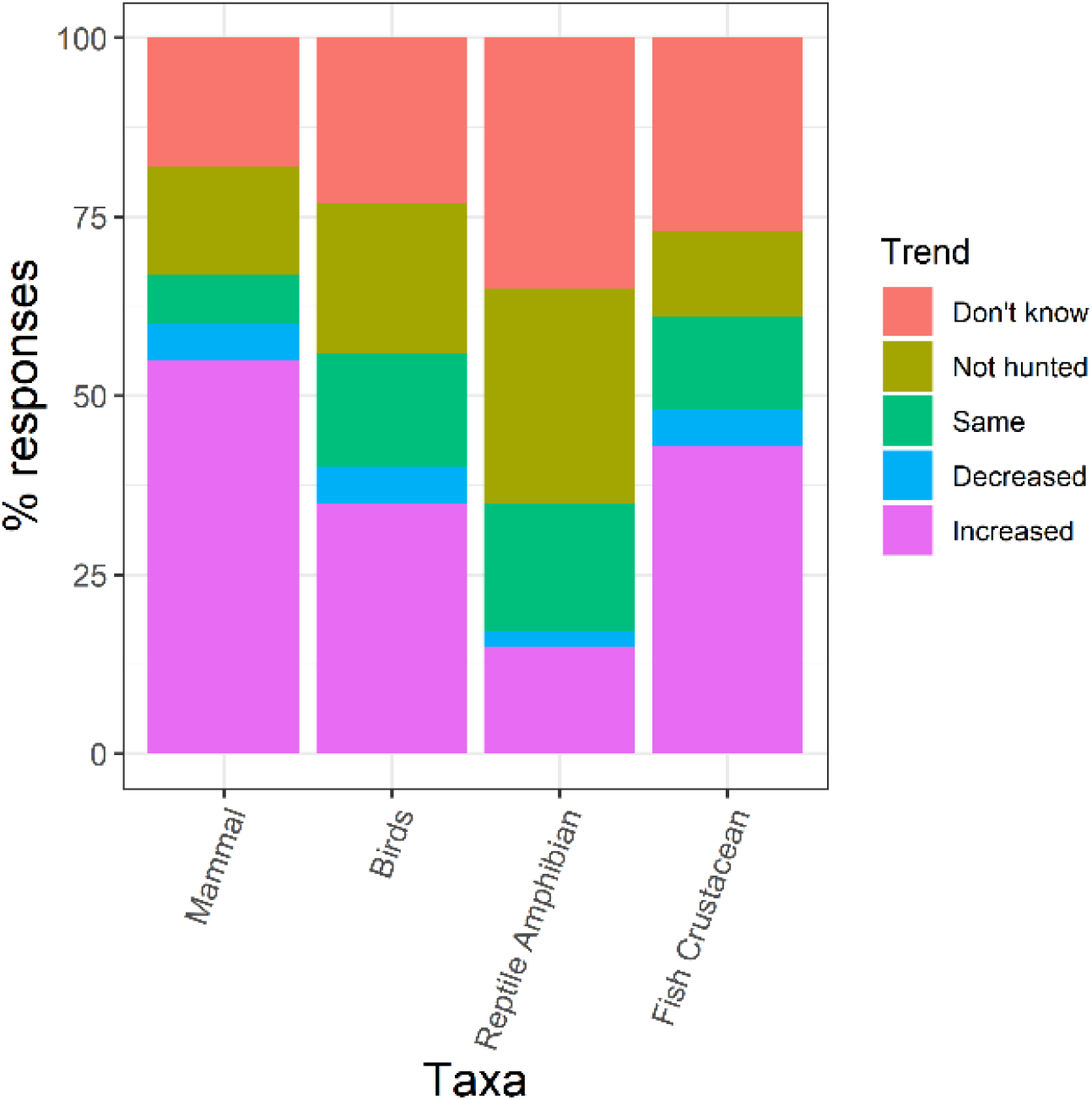
Change in hunting levels of different taxa during the lockdown based on answers by 99 respondents for each taxon (Q8, Supplementary Material 1).

Sixty-four percent (95% CI: 54% – 72%) stated that hunting during the lockdown was carried out by residents who were known to hunt regularly even before the lockdown, whilst 39% (95% CI: 29% – 49%) attributed hunting to residents who had lost employment due to the lockdown. Twenty percent (95% CI: 12% – 28%) reported hunting by individuals who moved back to this location during lockdown (Returnees), 17% (95% CI: 10% – 24%) reported hunting was done by mixed groups and 6 % (95% CI: 2% – 11%) by outsiders (Fig. 3a, Supplementary Material 2 Table 3). There was overlap in reported locations of hunting in Reserve Forests (43%; 95% CI: 34% – 53%), village revenue land (32%; 95% CI: 23% – 41%), Protected Areas (28%; 95% CI: 19% – 37%), private land (27%; 95% CI: 18% - 36%) and Territorial Forests (22%; 95% CI: 14% - 31%) (Fig. 3b, Supplementary Material 2 Table 4).

**Figure 3.**
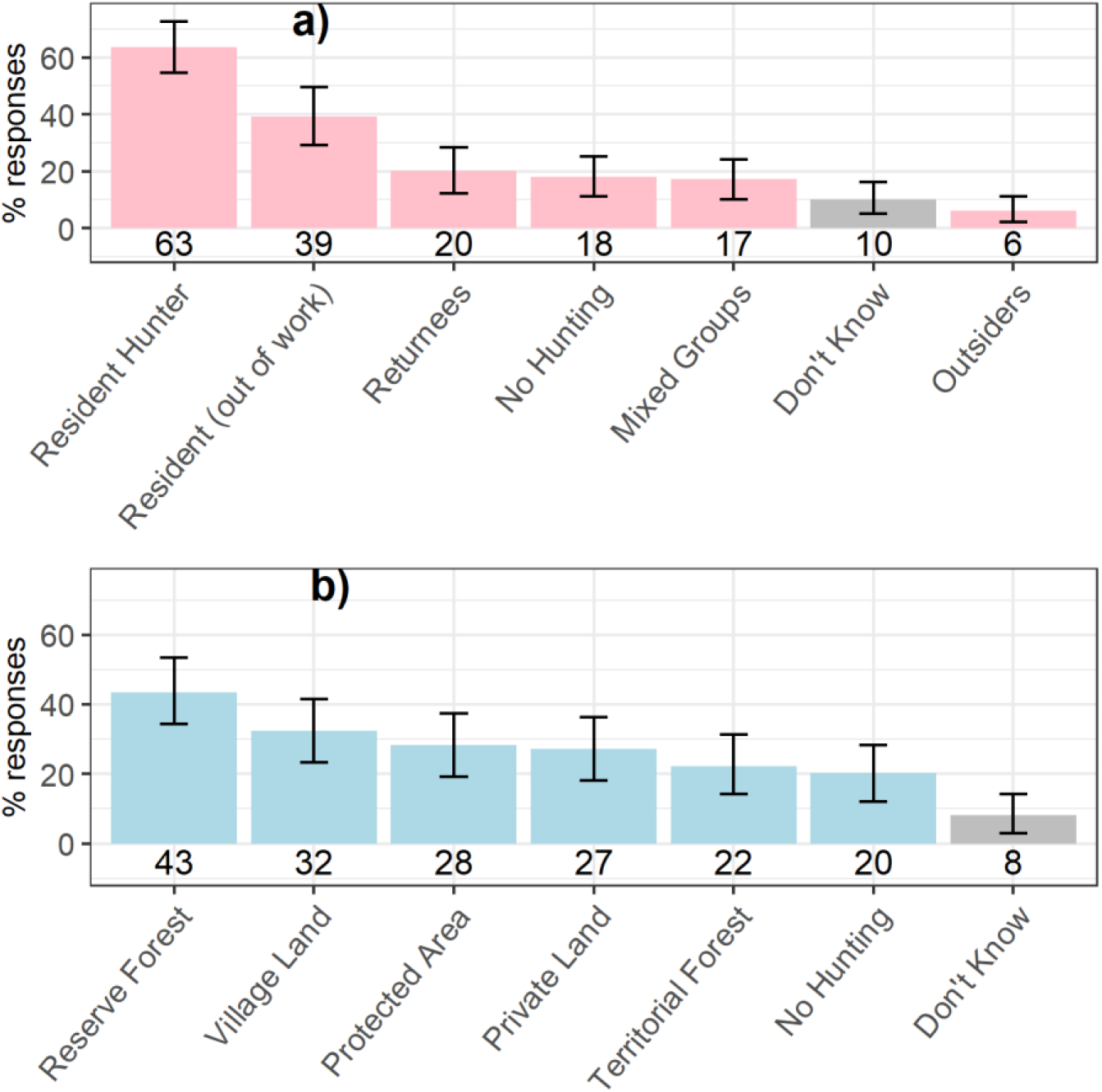
Bootstrapped mean and 95% CI of 99 respondents’ answers on (a) who was hunting (Q11, Supplementary Material 1) and (b) hunting location (Q9, Supplementary Material 1). Each respondent could choose more than one option for each question. The numbers below each bar are the number of respondents that choose that answer.

### 3.2 Questionnaire survey: motivations and factors

Over half the respondents (53%; 95% CI: 43% – 63%) felt that hunting for household consumption increased during the lockdown, 34% (95% CI: 25% – 43%) reported increased hunting for sport and recreation, followed by trade in local (14%; 95% CI: 8% – 21%) or outside (11%; 95% CI: 5% – 17%) markets. A further 12% (95% CI: 6% – 19%) reported increase in medicinal use (Fig. 4, Supplementary Material 2 Table 5).

**Figure 4.**
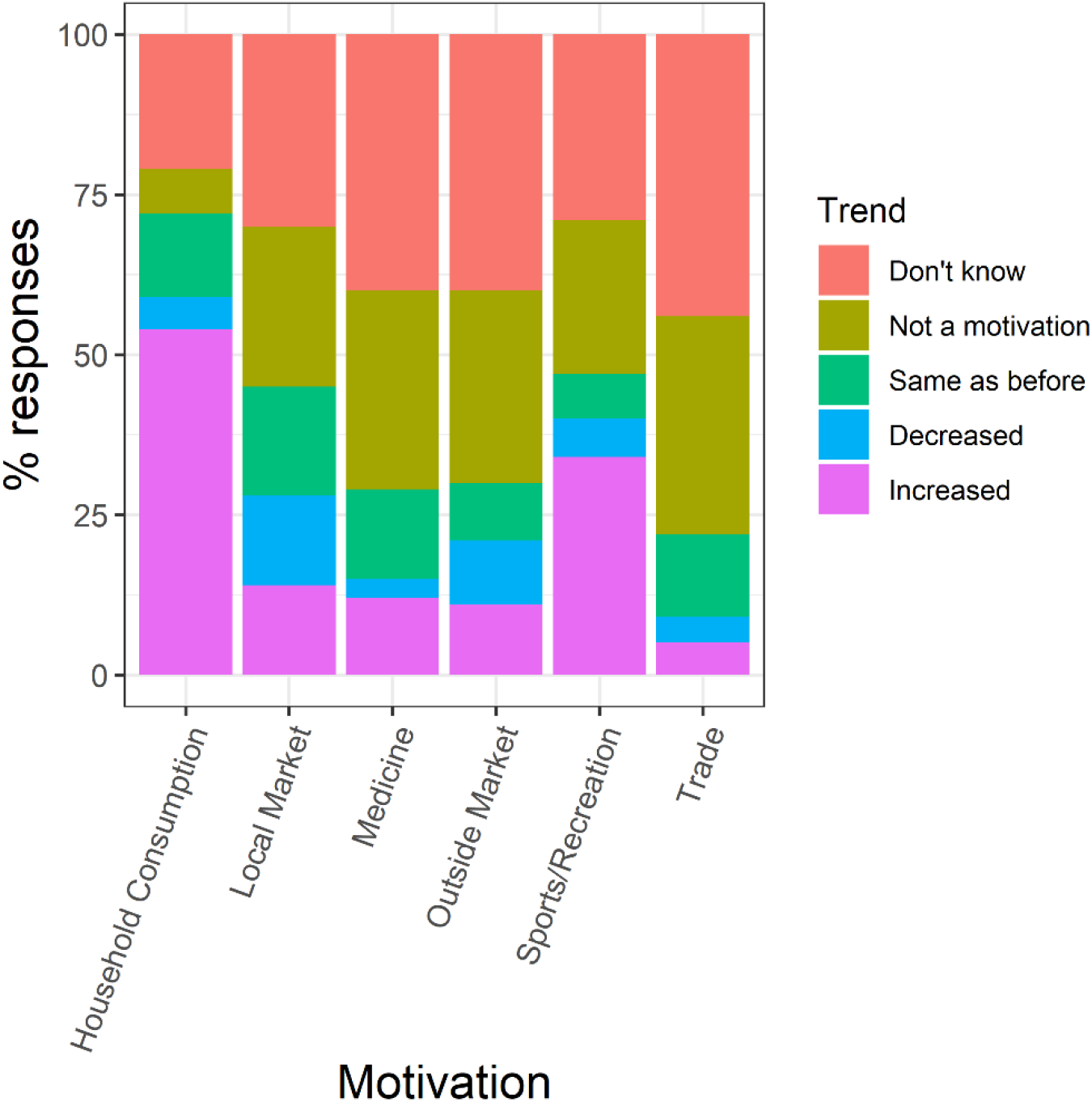
Change in motivations for hunting during the lockdown as answered by 99 respondents for each motivation. (Q10, Supplementary Material 1)

There was no association between perceived change in motivation and perceived change in hunting pressure of different taxa (Chi Sq test, X-squared = 6.8128, df = 15, p-value =0.9626 (Supplementary Material 2 Figure 1b), indicating no clear targeting of particular taxa during the lockdown.

There were overlapping factors associated with increase in hunting. More than one-third of the respondents (36%; 95% CI: 27% – 45%) felt that lack of enforcement during lockdown was a factor that resulted in the increase, 32% (95% CI: 23% – 41%) stated a disruption in food supplies and the same percentage 32% (95% CI: 23% – 41%) stated a need for recreation. Other factors were collapse of traditional seasonal occupations (24%; 95% CI: 16% - 33%), lack of incomes from tourism, handicrafts, and other local industries (21%; 95% CI: 14% - 29%), need for community bonding (18%; 95% CI: 11% – 26%) and need to supplement household income to sustain an influx of individuals from urban areas (7%; 95% CI: 2% - 12%) (Fig. 5, Supplementary Material 2 Table 6).

**Figure 5.**
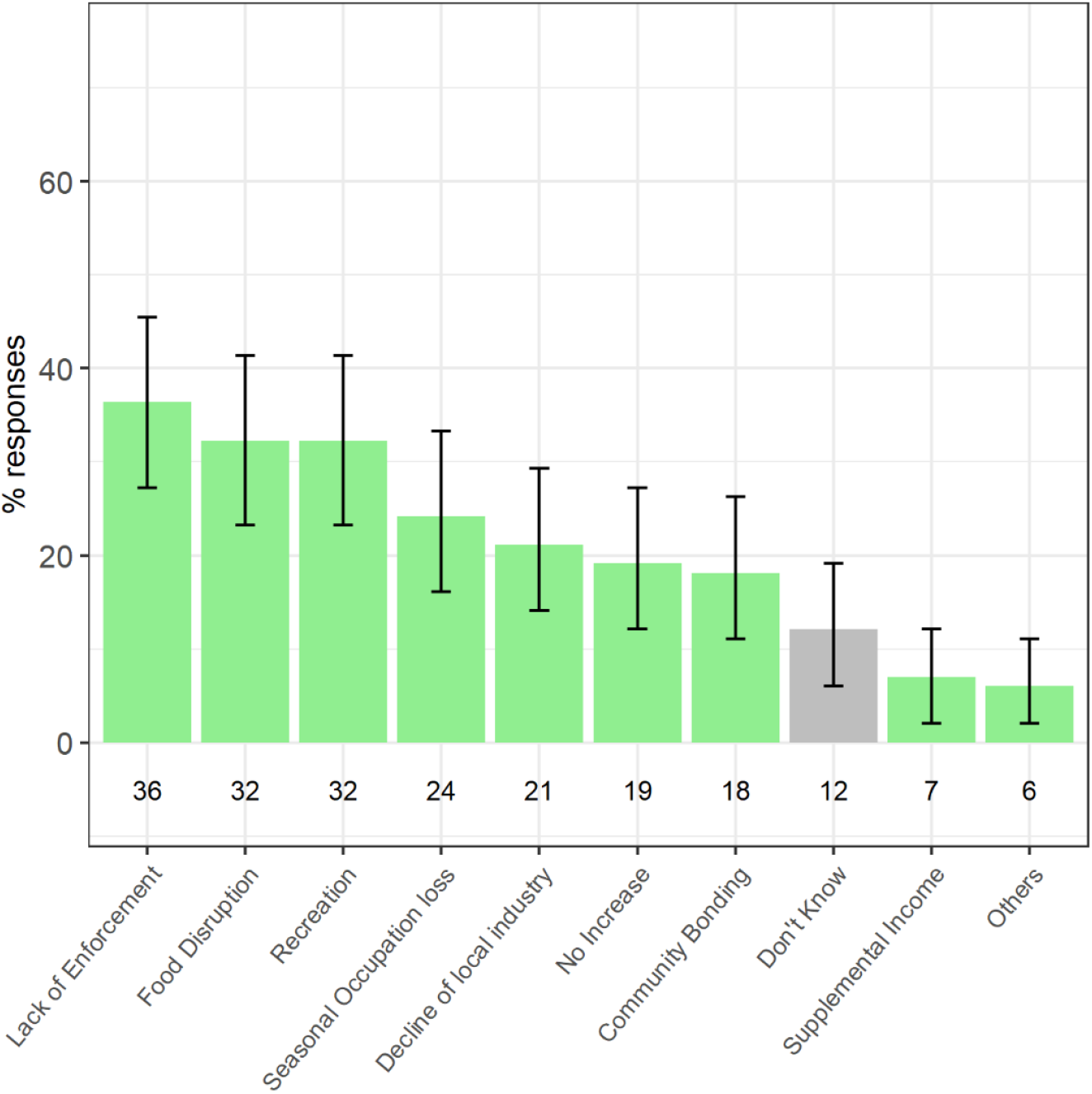
Bootstrapped mean and 95% CI of 99 respondents’ answers on what were the local factors associated with hunting (Q12, Supplementary Material 1). Each respondent could choose more than one option for each of the questions. The numbers below each bar are the number of respondents that choose that answer.

Our Chi-square test (X-squared = 13.784, df = 20, p-value = 0.8413) indicated that no single local factors were associated with change in motivation to hunt during lockdown (Supplementary Material 2 Fig. 1a).

### 3.3 Questionnaire survey: Counter-hunting strategies

At the same time, many respondents felt that enforcement action did not decline much across the different agencies with efforts remaining either the same for Forest Department (34%; 95% CI: 25% – 44%) and Police (29%; 95% CI: 20% – 38%), or increasing for Forest Department (20%; 95% CI: 12% – 28%) and Police (9%; 95% CI: 4% – 15%) (Supplementary Material 2 Table 7).

Respondents listed lack of staff strength (46%; 95% CI: 39% - 60%), lack of mobility (38%; 95% CI: 29% - 48%) and logistical constraints (38%; 95% CI: 29% - 48%) along with increased instances of hunting (36%; 95% CI: 27% - 45%) as major challenges for enforcement (Fig. 6, Supplementary Material 2 Table 8).

**Figure 6.**
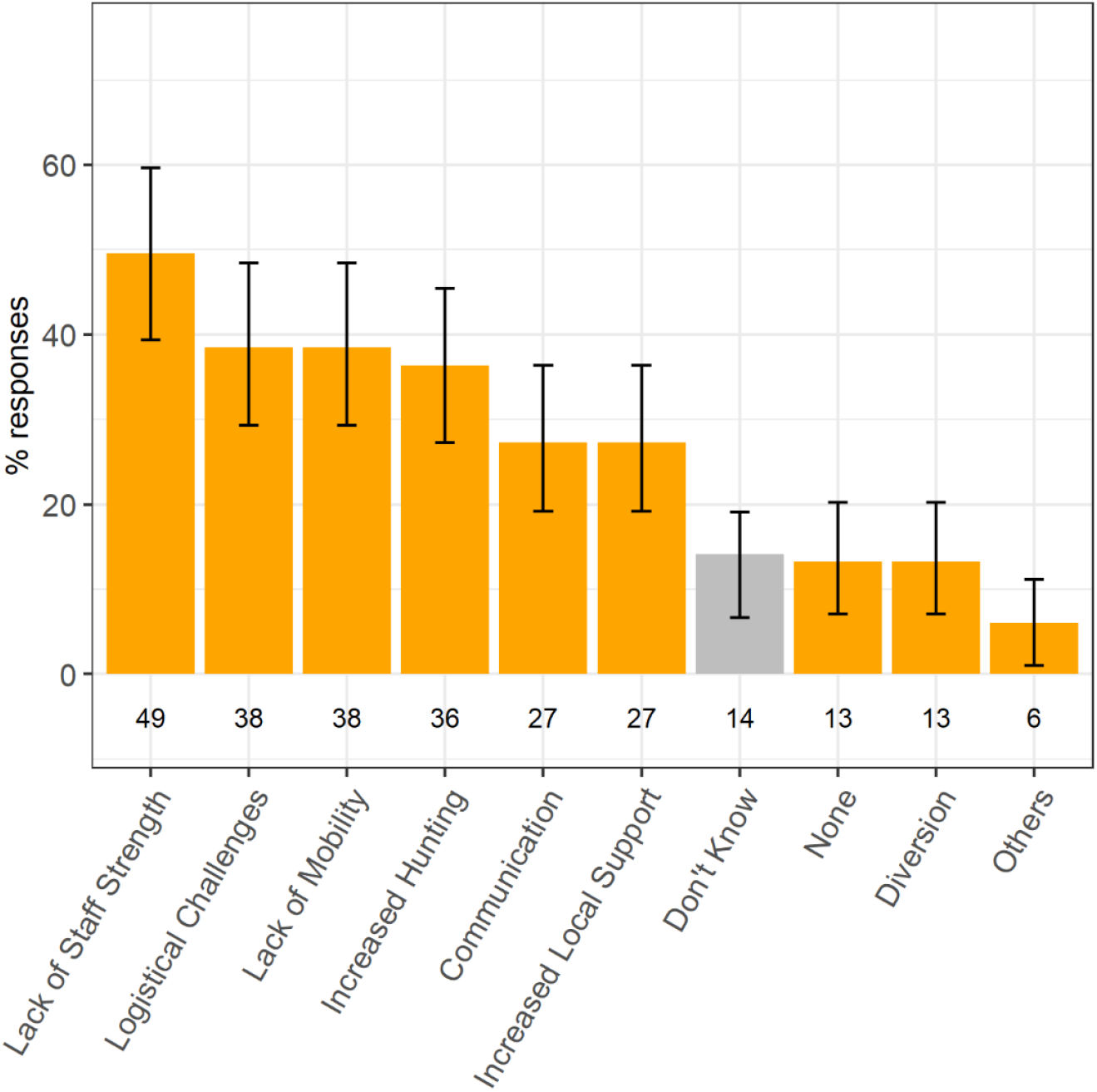
Bootstrapped mean and 95% CI of 99 respondents’ answers on what were the challenges faced by enforcement agencies during the lockdown (Q14, Supplementary Material 1). Each respondent could choose more than one option for each of the questions. The numbers below each bar are the number of respondents that choose that answer.

Information regarding strategies implemented by the administration and NGOs was sparse with a majority of respondents choosing ‘Don’t know’ for most options. However, nearly half (N= 47) respondents said that ‘provisioning of essential food supplies’ was implemented at their focal location, and of these 17% (95% CI: 10% – 25%) stated its efficacy at regulating hunting during the lockdown (Fig. 7, Supplementary Material Table 9)

**Figure 7.**
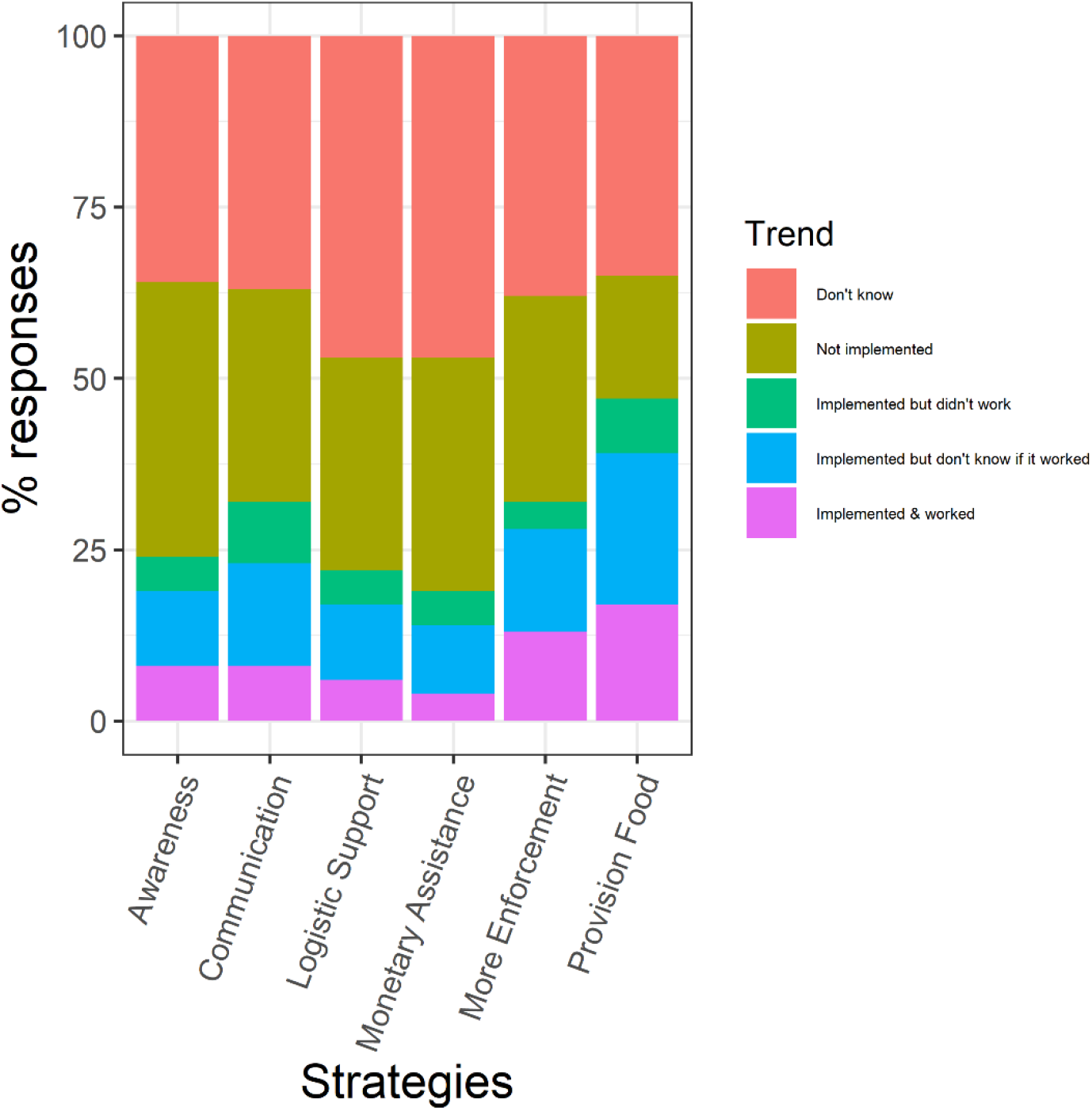
Strategies used to combat hunting during lockdown and their effectiveness as answered by 99 respondents for each strategy. (Q16, Supplementary Material 1)

### 3.4 Media analysis

Eighty-two percent of media statements (78 statements by 60 unique experts) suggested an increase in hunting during lockdown, 11% (10 unique expert statements) stated no change, 4% (4 statements by 2 unique experts) stated no hunting occurred, while 3% (3 unique expert statements) suggested a decrease in hunting occurred during lockdown. Increased hunting was recorded for mammals (19 statements), birds (6 statements) and reptiles/amphibians (5 statements), across sites in 12 different states (See Supplementary Material 3.1 for information regarding the media articles and the statements used to analyse the data).

In terms of motivation behind increased hunting, most statements (18 statements by 16 unique experts) indicated household consumption followed by sport and recreational (6 statements by 5 unique experts). Household consumption was primarily linked to food supply disruption (9 unique statements) and lack of income (5 unique statements), whilst sport and recreational hunting was linked to need for hobby during the lockdown (5 unique statements) (Supplementary Material 3.2 Table 2).

The media articles also had information on changes in enforcement by forest department (34 unique statements), community patrols (15 unique statements) and police department (3 unique) statements. The answers varied for each agency, as highlighted for the forest department, where in 10 unique statements suggested their enforcement against hunting remained the same, 18 unique statements suggested increased enforcement during lockdown, while 7 unique statements suggested a decreased during lockdown (Supplementary Material 3.2 Table 3). Broadly the qualitatively media analysis was similar to our questionnaire surveys.

## 4.0 Discussion

Our study suggests that many parts of India may have witnessed an increase in hunting during the COVID-19 lockdown. This increase seems to have been predominantly for household consumption, and to a lesser extent, for sport and recreation. Factors such as lower enforcement and disruption of food supply may have contributed to the perceived increase in hunting during the lockdown. Sale in local market and trade in animal body parts do not seem to have been affected significantly by the lockdown. Although, the increase in hunting during COVID-19 lockdowns has been reported by other studies (Aditya et al. 2021; Badola 2020; Manenti et al. 2020) our study provides unique insights into the motivations for this hunting and the effects of lockdowns.

We would like the readers to consider the following caveats: 1) data was collected from key respondents and media reports, not directly from hunters and therefore reflects perceptions rather than a real measure of hunting or motivations (Kahler and Gore 2015); 2) There were a number of “Don’t knows”, which might be attributed to low access to information during the lockdown, and hunting being understudied and a sensitive subject, especially within the Indian conservation scenario. We also acknowledge that the coarse scale of our data cannot reflect local nuances and tends towards oversimplification, for instance it is hard to distinguish illegal fishing from legal fishing (Oyanedel, Gelcich and Milner‐Gulland 2020).

We posit that one reason for the increase in hunting during the lockdown was the disruption of food supply chains. Shutting down of meat shops may have increased bushmeat demand, a possibility that has also been highlighted by eleven respondents in the open-ended section of our survey. Dietary habits vary dramatically across India. Some regions within the country are predominantly vegetarian and in other parts up to 90% of the households consume meat (Natrajan and Jacob 2018). The protein needs of people are met by inexpensive and easily accessible domestic protein options in most cases. However, in its absence it is possible that many would have turned to bushmeat consumption. Listing domestic meat shops as essential businesses along with grocery stores, especially in areas with high meat demand and during festivals that are marked with meat consumption becomes an important on ground consideration.

The lockdown also affected the food purchasing ability of millions of people across the country, especially those employed in the unorganized sector. We know that loss of jobs, especially by migrant workers, and the resulting food insecurity faced can have significant effects on use of natural resources (López-Feldman and Chávez, 2017; Tiwari and Joshi 2012). It is important for countries like India, that have numerous marginalized and poor groups, to consider measures to prevent widespread food insecurity during future lockdowns. Responses to our section related to strategies that worked to prevent hunting suggest that provisioning of essential food supplies may have worked to some extent, similar to recommendations from other experts (Ravallion 2020).

One-third of our survey respondents stated that there was an intensification in sport and recreational hunting during the lockdown, this was also corroborated by hunting videos from our media analyses (Shekar 2020). Although prevalence of recreational hunting, even within the Indian context has been acknowledged (Aiyadurai, Singh and Milner-Gulland 2010; Kaul, Jandrotia and McGowan 2004), our understanding of the value and motivation of recreational hunting and its effect on wildlife is still understudied (Chang et al. 2019).

Our attempt to understand the role of enforcement in preventing hunting during the lockdown was met with mixed results. Although over half our respondents felt there was no change or even an increase in the presence of enforcement agencies at their location, lack of enforcement was cited as a factor contributing to an increase in hunting by over one-third of the respondents. Some of this disparity can be explained by the fact that 16% of the respondents felt that increased instances of hunting during lockdown was a challenge for enforcement agencies. An increase in hunting linked to socioeconomic factors occur despite sustained enforcement (De Merode et al. 2007), implying that in addition to providing logistic support for enforcement, such as patrolling, there is a need to identify and address the socioeconomic drivers of hunting.

Another possible factor that may have played some role in increased non-compliance to hunting prohibitions might be resentment towards the government as has been suggested by one of the respondents who cited ‘anger against the government’ as a motivation. Studies have suggested that non-compliance with conservation regulations can stem from resentment towards the administration, especially enforcement agencies (Kahler and Gore 2015; Solomon, Gavin and Gore 2015).

Together the multitude of reasons related to hunting that unfold in this study highlight the significance of moving away from the notion of a singular mechanistic driver and to better cope with future socio-economic shocks that may result from pandemics, extreme climatic conditions, recessions, war and civil unrest.

## 5.0 Conclusion

For the foreseeable future, pandemic related restrictions and lockdowns are likely to have significant economic and social repercussions, which will create new challenges to effectively manage and conserve natural resources (Corlett et al. 2020). Even a year after the first COVID-19 lockdowns, various countries are seeing a series of renewed lockdowns in an effort to combat fresh waves of the pandemic (Reuters 2021; Richmond and Jordans 2020). It is imperative that in a COVID-19 world and beyond, alleviating shocks and setbacks (Carrington 2020; IPBES 2020;) will require developing rapid and novel response plans that include wildlife conservation and human-wellbeing around wildlife areas.

## Acknowledgments

We thank all the respondents who took time out to take our surveys. Without their effort our study would not be possible. We also thank everyone that helped circulate our survey through their networks. We appreciate discussions and comments provided by Anand Osuri. Additionally, a big thank you to the Ethics Committee of Nature Conservation Foundation (NCF) for scrutinizing the surveys and project for ethical appropriateness and giving very useful comments.

## Supplementary Material

Supplementary Material 1: Questionnaire Form

Supplementary Material 2: Questionnaire Results

Supplementary Material 3.1: Media Analysis - Raw Data

Supplementary Material 3.2: Media Analysis – Results

