## Supplementary Material 1 for "Key informant perceptions on wildlife hunting in India during the COVID-19 lockdown"

### How has the nationwide lock-down affected wildlife hunting in India?

Hello, we hope you are keeping healthy and safe during this unprecedented crisis.

We are conducting an anonymous online survey of key-informants, wildlife experts, managers and enthusiasts (over the age of 18) to understand the impact of nationwide COVID-19 lock-down on illegal hunting and forest management in India. We are doing this by comparing patterns of hunting between the lock-down period (specifically, 25 March to 4 May 2020, corresponding to the first two lock-down phases), and the pre-lock-down period (23 Jan-24 March 2020). We request you to volunteer 10-15 minutes to share your knowledge on this matter by responding to the questions below. If you wish to report for more than one location please fill a fresh survey form for each location.

Please be assured that your confidentiality is our priority. We are not recording any personal identification information in this survey, and neither we nor anyone else that views the data or results will be able to trace responses to individual respondents. Our survey questions have been reviewed and approved by the Research Ethics Committee of Nature Conservation Foundation.

Please share this survey with others who you think will be in a position to contribute to this study. The survey will remain open till 20 May 2020, but we urge you to respond soon. Feel free to reach out to us if you have any questions.

Note: By submitting this form you are consenting to your anonymous responses being used for research and conservation projects, and to the data being archived in a public repository. You will not receive personal (direct) or monetary benefits from taking part in this research study.

Thank you,

Uttara Mendiratta,
Nirmal Kulkarni,
Nandini Velho, &
Kulbhushan Suryawanshi

*Required

1. 1) Which one of these best describes your professional role *

*Tick all that apply.*

- [ ] Government agency
- [ ] University or academic organisation
- [ ] Non-government organisation (NGO) – research
- [ ] NGO – non research
- [ ] Tourism and allied businesses
- [ ] Journalism/photography
- [ ] Agriculture/plantations

Other: [ ] ______________________

2. 2) What is your age? *

*Mark only one oval.*

- ( ) 18-24
- ( ) 25-34
- ( ) 35-44
- ( ) 45-54
- ( ) 55-64
- ( ) 65-74
- ( ) >75
- ( ) Prefer not to say

3. 3) What is your gender? *

*Mark only one oval.*

- ( ) Female
- ( ) Male
- ( ) Other
- ( ) Prefer not to say

4. 4.1) State/Union Territory & District of the location that you are reporting about *

*Mark only one oval.*

- Andaman & Nicobar Island, Nicobar
- Andaman & Nicobar Island, South Andaman
- Andaman & Nicobar Island, North & Middle Andaman
- Andhra Pradesh, Adilabad
- Andhra Pradesh, Anantapur
- Andhra Pradesh, Chittoor
- Andhra Pradesh, Y.s.r.
- Andhra Pradesh, East Godavari
- Andhra Pradesh, Guntur
- Andhra Pradesh, Hyderabad
- Andhra Pradesh, Karimnagar
- Andhra Pradesh, Khammam
- Andhra Pradesh, Krishna
- Andhra Pradesh, Kurnool
- Andhra Pradesh, Mahbubnagar
- Andhra Pradesh, Medak
- Andhra Pradesh, Nalgonda
- Andhra Pradesh, Sri Potti Sriramulu Nellore
- Andhra Pradesh, Nizamabad
- Andhra Pradesh, Prakasam
- Andhra Pradesh, Rangareddy
- Andhra Pradesh, Srikakulam
- Andhra Pradesh, Visakhapatnam
- Andhra Pradesh, Vizianagaram
- Andhra Pradesh, Warangal
- Andhra Pradesh, West Godavari
- Arunanchal Pradesh, Anjaw
- Arunanchal Pradesh, Changlang
- Arunanchal Pradesh, Dibang Valley
- Arunanchal Pradesh, East Kameng
- Arunanchal Pradesh, East Siang
- Arunanchal Pradesh, Kurung Kumey

- Arunanchal Pradesh, Lohit
- Arunanchal Pradesh, Lower Dibang Valley
- Arunanchal Pradesh, Lower Subansiri
- Arunanchal Pradesh, Papum Pare
- Arunanchal Pradesh, Tawang
- Arunanchal Pradesh, Tirap
- Arunanchal Pradesh, Upper Siang
- Arunanchal Pradesh, Upper Subansiri
- Arunanchal Pradesh, West Kameng
- Arunanchal Pradesh, West Siang
- Assam, Baksa
- Assam, Barpeta
- Assam, Bongaigaon
- Assam, Cachar
- Assam, Chirang
- Assam, Darrang
- Assam, Dhemaji
- Assam, Dhubri
- Assam, Dibrugarh
- Assam, Goalpara
- Assam, Golaghat
- Assam, Hailakandi
- Assam, Jorhat
- Assam, Kamrup
- Assam, Kamrup Metropolitan
- Assam, Karbi Anglong
- Assam, Karimganj
- Assam, Kokrajhar
- Assam, Lakhimpur
- Assam, Marigaon
- Assam, Nagaon
- Assam, Nalbari
- Assam, Dima Hasao
- Assam, Sivasagar
- Assam, Sonitpur

- Assam, Tinsukia
- Assam, Udalguri
- Bihar, Araria
- Bihar, Aurangabad
- Bihar, Banka
- Bihar, Begusarai
- Bihar, Bhagalpur
- Bihar, Bhojpur
- Bihar, Buxar
- Bihar, Darbhanga
- Bihar, Gaya
- Bihar, Gopalganj
- Bihar, Jamui
- Bihar, Kaimur (bhabua)
- Bihar, Katihar
- Bihar, Khagaria
- Bihar, Kishanganj
- Bihar, Lakhisarai
- Bihar, Madhepura
- Bihar, Madhubani
- Bihar, Munger
- Bihar, Muzaffarpur
- Bihar, Nalanda
- Bihar, Nawada
- Bihar, Pashchim Champaran
- Bihar, Patna
- Bihar, Purba Champaran
- Bihar, Purnia
- Bihar, Rohtas
- Bihar, Saharsa
- Bihar, Samastipur
- Bihar, Saran (chhapra)
- Bihar, Sheikhpura
- Bihar, Sheohar
- Bihar, Sitamarhi

- Bihar, Siwan
- Bihar, Supaul
- Bihar, Vaishali
- Bihar, Arwal
- Bihar, Jehanabad
- Chandigarh, Chandigarh
- Chhattisgarh, Bastar
- Chhattisgarh, Bijapur
- Chhattisgarh, Bilaspur
- Chhattisgarh, Dakshin Bastar Dantewada
- Chhattisgarh, Dhamtari
- Chhattisgarh, Durg
- Chhattisgarh, Janjgir-champa
- Chhattisgarh, Jashpur
- Chhattisgarh, Uttar Bastar Kanker
- Chhattisgarh, Korba
- Chhattisgarh, Koriya
- Chhattisgarh, Mahasamund
- Chhattisgarh, Narayanpur
- Chhattisgarh, Raigarh
- Chhattisgarh, Raipur
- Chhattisgarh, Rajnandgaon
- Chhattisgarh, Surguja
- Chhattisgarh, Kabeerdham
- Dadara & Nagar Havelli, Dadra & Nagar Haveli
- Daman & Diu, Daman
- Daman & Diu, Diu
- Goa, North Goa
- Goa, South Goa
- Gujarat, Ahmadabad
- Gujarat, Amreli
- Gujarat, Anand
- Gujarat, Banas Kantha
- Gujarat, Bharuch
- Gujarat, Bhavnagar

- Gujarat, Dohad
- Gujarat, Gandhinagar
- Gujarat, Jamnagar
- Gujarat, Junagadh
- Gujarat, Kachchh
- Gujarat, Kheda
- Gujarat, Mahesana
- Gujarat, Narmada
- Gujarat, Navsari
- Gujarat, Panch Mahals
- Gujarat, Patan
- Gujarat, Porbandar
- Gujarat, Rajkot
- Gujarat, Sabar Kantha
- Gujarat, Surat
- Gujarat, Surendranagar
- Gujarat, The Dangs
- Gujarat, Vadodara
- Gujarat, Valsad
- Gujarat, Tapi
- Haryana, Ambala
- Haryana, Bhiwani
- Haryana, Faridabad
- Haryana, Fatehabad
- Haryana, Gurgaon
- Haryana, Hisar
- Haryana, Jhajjar
- Haryana, Kaithal
- Haryana, Karnal
- Haryana, Kurukshetra
- Haryana, Mahendragarh
- Haryana, Mewat
- Haryana, Palwal
- Haryana, Panchkula
- Haryana, Panipat

- Haryana, Rewari
- Haryana, Rohtak
- Haryana, Sirsa
- Haryana, Sonipat
- Haryana, Yamunanagar
- Haryana, Jind
- Himachal Pradesh, Bilaspur
- Himachal Pradesh, Chamba
- Himachal Pradesh, Hamirpur
- Himachal Pradesh, Kangra
- Himachal Pradesh, Kinnaur
- Himachal Pradesh, Kullu
- Himachal Pradesh, Lahul & Spiti
- Himachal Pradesh, Mandi
- Himachal Pradesh, Shimla
- Himachal Pradesh, Sirmaur
- Himachal Pradesh, Solan
- Himachal Pradesh, Una
- Jammu & Kashmir, Anantnag
- Jammu & Kashmir, Badgam
- Jammu & Kashmir, Bandipore
- Jammu & Kashmir, Baramula
- Jammu & Kashmir, Data Not Available
- Jammu & Kashmir, Doda
- Jammu & Kashmir, Ganderbal
- Jammu & Kashmir, Jammu
- Jammu & Kashmir, Kargil
- Jammu & Kashmir, Kathua
- Jammu & Kashmir, Kishtwar
- Jammu & Kashmir, Kulgam
- Jammu & Kashmir, Kupwara
- Jammu & Kashmir, Leh (ladakh)
- Jammu & Kashmir, Pulwama
- Jammu & Kashmir, Punch
- Jammu & Kashmir, Rajouri

- Jammu & Kashmir, Ramban
- Jammu & Kashmir, Reasi
- Jammu & Kashmir, Samba
- Jammu & Kashmir, Shupiyan
- Jammu & Kashmir, Srinagar
- Jammu & Kashmir, Udhampur
- Jharkhand, Bokaro
- Jharkhand, Chatra
- Jharkhand, Deoghar
- Jharkhand, Dhanbad
- Jharkhand, Dumka
- Jharkhand, Garhwa
- Jharkhand, Giridih
- Jharkhand, Godda
- Jharkhand, Gumla
- Jharkhand, Hazaribagh
- Jharkhand, Jamtara
- Jharkhand, Khunti
- Jharkhand, Kodarma
- Jharkhand, Latehar
- Jharkhand, Lohardaga
- Jharkhand, Pakur
- Jharkhand, Palamu
- Jharkhand, Pashchimi Singhbhum
- Jharkhand, Purbi Singhbhum
- Jharkhand, Ramgarh
- Jharkhand, Ranchi
- Jharkhand, Sahibganj
- Jharkhand, Saraikela-kharsawan
- Jharkhand, Simdega
- Karnataka, Bagalkot
- Karnataka, Bangalore Rural
- Karnataka, Bangalore
- Karnataka, Belgaum
- Karnataka, Bellary

- Karnataka, Bidar
- Karnataka, Bijapur
- Karnataka, Chamrajnagar
- Karnataka, Chikkaballapura
- Karnataka, Chikmagalur
- Karnataka, Chitradurga
- Karnataka, Dakshina Kannada
- Karnataka, Davanagere
- Karnataka, Dharwad
- Karnataka, Gadag
- Karnataka, Gulbarga
- Karnataka, Hassan
- Karnataka, Haveri
- Karnataka, Kodagu
- Karnataka, Kolar
- Karnataka, Koppal
- Karnataka, Mandya
- Karnataka, Mysore
- Karnataka, Raichur
- Karnataka, Ramanagara
- Karnataka, Shimoga
- Karnataka, Tumkur
- Karnataka, Udupi
- Karnataka, Uttara Kannada
- Karnataka, Yadgir
- Kerala, Alappuzha
- Kerala, Ernakulam
- Kerala, Idukki
- Kerala, Kannur
- Kerala, Kasaragod
- Kerala, Kollam
- Kerala, Kottayam
- Kerala, Kozhikode
- Kerala, Malappuram
- Kerala, Palakkad

- Kerala, Pathanamthitta
- Kerala, Thiruvananthapuram
- Kerala, Thrissur
- Kerala, Wayanad
- Lakshadweep, Lakshadweep
- Madhya Pradesh, Alirajpur
- Madhya Pradesh, Anuppur
- Madhya Pradesh, Ashoknagar
- Madhya Pradesh, Balaghat
- Madhya Pradesh, Barwani
- Madhya Pradesh, Betul
- Madhya Pradesh, Bhind
- Madhya Pradesh, Bhopal
- Madhya Pradesh, Burhanpur
- Madhya Pradesh, Chhatarpur
- Madhya Pradesh, Chhindwara
- Madhya Pradesh, Damoh
- Madhya Pradesh, Datia
- Madhya Pradesh, Dewas
- Madhya Pradesh, Dhar
- Madhya Pradesh, Dindori
- Madhya Pradesh, East Nimar
- Madhya Pradesh, Guna
- Madhya Pradesh, Gwalior
- Madhya Pradesh, Harda
- Madhya Pradesh, Hoshangabad
- Madhya Pradesh, Indore
- Madhya Pradesh, Jabalpur
- Madhya Pradesh, Jhabua
- Madhya Pradesh, Katni
- Madhya Pradesh, Mandla
- Madhya Pradesh, Mandsaur
- Madhya Pradesh, Morena
- Madhya Pradesh, Narsimhapur
- Madhya Pradesh, Neemuch

- Madhya Pradesh, Panna
- Madhya Pradesh, Raisen
- Madhya Pradesh, Rajgarh
- Madhya Pradesh, Ratlam
- Madhya Pradesh, Rewa
- Madhya Pradesh, Sagar
- Madhya Pradesh, Satna
- Madhya Pradesh, Sehore
- Madhya Pradesh, Seoni
- Madhya Pradesh, Shahdol
- Madhya Pradesh, Shajapur
- Madhya Pradesh, Sheopur
- Madhya Pradesh, Shivpuri
- Madhya Pradesh, Sidhi
- Madhya Pradesh, Singrauli
- Madhya Pradesh, Tikamgarh
- Madhya Pradesh, Ujjain
- Madhya Pradesh, Umaria
- Madhya Pradesh, Vidisha
- Madhya Pradesh, West Nimar
- Maharashtra, Ahmadnagar
- Maharashtra, Akola
- Maharashtra, Amravati
- Maharashtra, Aurangabad
- Maharashtra, Bhandara
- Maharashtra, Bid
- Maharashtra, Buldana
- Maharashtra, Chandrapur
- Maharashtra, Dhule
- Maharashtra, Garhchiroli
- Maharashtra, Gondiya
- Maharashtra, Hingoli
- Maharashtra, Jalgaon
- Maharashtra, Jalna
- Maharashtra, Kolhapur

- Maharashtra, Latur
- Maharashtra, Mumbai
- Maharashtra, Mumbai Suburban
- Maharashtra, Nagpur
- Maharashtra, Nanded
- Maharashtra, Nandurbar
- Maharashtra, Nashik
- Maharashtra, Osmanabad
- Maharashtra, Parbhani
- Maharashtra, Pune
- Maharashtra, Raigarh
- Maharashtra, Ratnagiri
- Maharashtra, Sangli
- Maharashtra, Satara
- Maharashtra, Sindhudurg
- Maharashtra, Solapur
- Maharashtra, Thane
- Maharashtra, Wardha
- Maharashtra, Washim
- Maharashtra, Yavatmal
- Manipur, Bishnupur
- Manipur, Chandel
- Manipur, Churachandpur
- Manipur, Imphal East
- Manipur, Imphal West
- Manipur, Senapati
- Manipur, Tamenglong
- Manipur, Thoubal
- Manipur, Ukhrul
- Meghalaya, East Garo Hills
- Meghalaya, East Khasi Hills
- Meghalaya, Jaintia Hills
- Meghalaya, Ri Bhoi
- Meghalaya, South Garo Hills
- Meghalaya, West Garo Hills

- Meghalaya, West Khasi Hills
- Mizoram, Aizawl
- Mizoram, Champhai
- Mizoram, Kolasib
- Mizoram, Lawangtlai
- Mizoram, Lunglei
- Mizoram, Mamit
- Mizoram, Saiha
- Mizoram, Serchhip
- Nagaland, Dimapur
- Nagaland, Kiphire
- Nagaland, Kohima
- Nagaland, Longleng
- Nagaland, Mokokchung
- Nagaland, Mon
- Nagaland, Peren
- Nagaland, Phek
- Nagaland, Tuensang
- Nagaland, Wokha
- Nagaland, Zunheboto
- NCT of Delhi, Central
- NCT of Delhi, East
- NCT of Delhi, New Delhi
- NCT of Delhi, North
- NCT of Delhi, North East
- NCT of Delhi, North West
- NCT of Delhi, South
- NCT of Delhi, South West
- NCT of Delhi, West
- Odisha, Anugul
- Odisha, Balangir
- Odisha, Baleshwar
- Odisha, Bargarh
- Odisha, Bauda
- Odisha, Bhadrak

- Odisha, Cuttack
- Odisha, Debagarh
- Odisha, Dhenkanal
- Odisha, Gajapati
- Odisha, Ganjam
- Odisha, Jagatsinghapur
- Odisha, Jajapur
- Odisha, Jharsuguda
- Odisha, Kalahandi
- Odisha, Kandhamal
- Odisha, Kendrapara
- Odisha, Kendujhar
- Odisha, Khordha
- Odisha, Koraput
- Odisha, Malkangiri
- Odisha, Mayurbhanj
- Odisha, Nabarangapur
- Odisha, Nayagarh
- Odisha, Nuapada
- Odisha, Puri
- Odisha, Rayagada
- Odisha, Sambalpur
- Odisha, Subarnapur
- Odisha, Sundargarh
- Puducherry, Mahe
- Puducherry, Karaikal
- Puducherry, Puducherry
- Puducherry, Yanam
- Punjab, Amritsar
- Punjab, Barnala
- Punjab, Bathinda
- Punjab, Faridkot
- Punjab, Fatehgarh Sahib
- Punjab, Firozpur
- Punjab, Gurdaspur

- Punjab, Hoshiarpur
- Punjab, Jalandhar
- Punjab, Kapurthala
- Punjab, Ludhiana
- Punjab, Mansa
- Punjab, Moga
- Punjab, Muktsar
- Punjab, Patiala
- Punjab, Rupnagar
- Punjab, Sahibzada Ajit Singh Nagar
- Punjab, Sangrur
- Punjab, Shahid Bhagat Singh Nagar
- Punjab, Tarn Taran
- Rajasthan, Ajmer
- Rajasthan, Alwar
- Rajasthan, Banswara
- Rajasthan, Baran
- Rajasthan, Barmer
- Rajasthan, Bharatpur
- Rajasthan, Bhilwara
- Rajasthan, Bikaner
- Rajasthan, Bundi
- Rajasthan, Chittaurgarh
- Rajasthan, Churu
- Rajasthan, Dhaulpur
- Rajasthan, Dungarpur
- Rajasthan, Ganganagar
- Rajasthan, Hanumangarh
- Rajasthan, Jaipur
- Rajasthan, Jaisalmer
- Rajasthan, Jalor
- Rajasthan, Jhalawar
- Rajasthan, Jhunjhunun
- Rajasthan, Jodhpur
- Rajasthan, Kota

- Rajasthan, Nagaur
- Rajasthan, Pali
- Rajasthan, Pratapgarh
- Rajasthan, Rajsamand
- Rajasthan, Sikar
- Rajasthan, Sirohi
- Rajasthan, Tonk
- Rajasthan, Udaipur
- Rajasthan, Dausa
- Rajasthan, Karauli
- Rajasthan, Sawai Madhopur
- Sikkim, East
- Sikkim, North
- Sikkim, South
- Sikkim, West
- Tamil Nadu, Ariyalur
- Tamil Nadu, Chennai
- Tamil Nadu, Coimbatore
- Tamil Nadu, Cuddalore
- Tamil Nadu, Dharmapuri
- Tamil Nadu, Dindigul
- Tamil Nadu, Erode
- Tamil Nadu, Kancheepuram
- Tamil Nadu, Kanniyakumari
- Tamil Nadu, Karur
- Tamil Nadu, Krishnagiri
- Tamil Nadu, Madurai
- Tamil Nadu, Namakkal
- Tamil Nadu, Perambalur
- Tamil Nadu, Pudukkottai
- Tamil Nadu, Ramanathapuram
- Tamil Nadu, Salem
- Tamil Nadu, Sivaganga
- Tamil Nadu, Thanjavur
- Tamil Nadu, The Nilgiris

- Tamil Nadu, Theni
- Tamil Nadu, Thiruvallur
- Tamil Nadu, Thiruvarur
- Tamil Nadu, Thoothukkudi
- Tamil Nadu, Tiruchirappalli
- Tamil Nadu, Tirunelveli
- Tamil Nadu, Tiruppur
- Tamil Nadu, Tiruvannamalai
- Tamil Nadu, Vellore
- Tamil Nadu, Viluppuram
- Tamil Nadu, Virudunagar
- Tamil Nadu, Nagappattinam
- Tripura, Dhalai
- Tripura, North Tripura
- Tripura, South Tripura
- Tripura, West Tripura
- Uttar Pradesh, Agra
- Uttar Pradesh, Aligarh
- Uttar Pradesh, Allahabad
- Uttar Pradesh, Ambedkar Nagar
- Uttar Pradesh, Auraiya
- Uttar Pradesh, Azamgarh
- Uttar Pradesh, Baghpat
- Uttar Pradesh, Bahraich
- Uttar Pradesh, Ballia
- Uttar Pradesh, Balrampur
- Uttar Pradesh, Banda
- Uttar Pradesh, Bara Banki
- Uttar Pradesh, Bareilly
- Uttar Pradesh, Basti
- Uttar Pradesh, Bijnor
- Uttar Pradesh, Budaun
- Uttar Pradesh, Bulandshahr
- Uttar Pradesh, Chandauli
- Uttar Pradesh, Chitrakoot

- Uttar Pradesh, Deoria
- Uttar Pradesh, Etah
- Uttar Pradesh, Etawah
- Uttar Pradesh, Faizabad
- Uttar Pradesh, Farrukhabad
- Uttar Pradesh, Fatehpur
- Uttar Pradesh, Firozabad
- Uttar Pradesh, Gautam Buddha Nagar
- Uttar Pradesh, Ghaziabad
- Uttar Pradesh, Ghazipur
- Uttar Pradesh, Gonda
- Uttar Pradesh, Gorakhpur
- Uttar Pradesh, Hamirpur
- Uttar Pradesh, Hardoi
- Uttar Pradesh, Mahamaya Nagar
- Uttar Pradesh, Jalaun
- Uttar Pradesh, Jaunpur
- Uttar Pradesh, Jhansi
- Uttar Pradesh, Jyotiba Phule Nagar
- Uttar Pradesh, Kannauj
- Uttar Pradesh, Kanpur Dehat
- Uttar Pradesh, Kanpur Nagar
- Uttar Pradesh, Kansiram Nagar
- Uttar Pradesh, Kaushambi
- Uttar Pradesh, Kheri
- Uttar Pradesh, Kushinagar
- Uttar Pradesh, Lalitpur
- Uttar Pradesh, Lucknow
- Uttar Pradesh, Maharajganj
- Uttar Pradesh, Mahoba
- Uttar Pradesh, Mainpuri
- Uttar Pradesh, Mathura
- Uttar Pradesh, Mau
- Uttar Pradesh, Meerut
- Uttar Pradesh, Mirzapur

- Uttar Pradesh, Moradabad
- Uttar Pradesh, Muzaffarnagar
- Uttar Pradesh, Pilibhit
- Uttar Pradesh, Pratapgarh
- Uttar Pradesh, Rae Bareli
- Uttar Pradesh, Rampur
- Uttar Pradesh, Saharanpur
- Uttar Pradesh, Sant Kabir Nagar
- Uttar Pradesh, Sant Ravi Das Nagar(bhadohi)
- Uttar Pradesh, Shahjahanpur
- Uttar Pradesh, Shrawasti
- Uttar Pradesh, Siddharth Nagar
- Uttar Pradesh, Sitapur
- Uttar Pradesh, Sonbhadra
- Uttar Pradesh, Sultanpur
- Uttar Pradesh, Unnao
- Uttar Pradesh, Varanasi
- Uttarakhand, Almora
- Uttarakhand, Bageshwar
- Uttarakhand, Chamoli
- Uttarakhand, Champawat
- Uttarakhand, Dehradun
- Uttarakhand, Garhwal
- Uttarakhand, Hardwar
- Uttarakhand, Nainital
- Uttarakhand, Pithoragarh
- Uttarakhand, Rudraprayag
- Uttarakhand, Tehri Garhwal
- Uttarakhand, Udham Singh Nagar
- Uttarakhand, Uttarkashi
- West Bengal, Bankura
- West Bengal, Barddhaman
- West Bengal, Birbhum
- West Bengal, Dakshin Dinajpur
- West Bengal, Darjiling

- West Bengal, Haora
- West Bengal, Hugli
- West Bengal, Jalpaiguri
- West Bengal, Koch Bihar
- West Bengal, Kolkata
- West Bengal, Maldah
- West Bengal, Murshidabad
- West Bengal, Nadia
- West Bengal, North 24 Parganas
- West Bengal, Pashchim Medinipur
- West Bengal, Purba Medinipur
- West Bengal, Puruliya
- West Bengal, South 24 Parganas
- West Bengal, Uttar Dinajpur
- OTHER

5. 4.2) If your State/UT, District name did not appear on the list above please choose 'Other' and give the details here -

_______________________________

6. 5) What is your source of information for this location? *

*Mark only one oval.*

- DIRECT (you were based at the location during lock-down)
- INDIRECT (you had direct contact with colleagues/assistants/collaborators at location during lock-down)
- BOTH (DIRECT & INDIRECT)

7. 6) What were the prevalent indicators of hunting at this location during the COVID-19 lock-down (phases 1&2 only – 25 March- 3 May)? (Tick all applicable options) *

*Tick all that apply.*

- [ ] No hunting
- [ ] Direct sighting or knowledge of hunting
- [ ] Snares/traps
- [ ] Gunshots heard
- [ ] Illegal fishing
- [ ] Wild animals sold in local markets
- [ ] Hunting parties with traps/guns/dogs
- [ ] Burning grasslands to hunt
- [ ] Action by local authorities in response to a hunting incident

Other: [ ] ____________________

8. 7) Did hunting levels change during the first two phases of the lock-down (25 March - 3 May) compared to the two months before the lock-down (23 Jan - 24 March) at this location? *

*Mark only one oval.*

- ( ) Drastic increase
- ( ) Some increase
- ( ) No change
- ( ) Some decrease
- ( ) Drastic decrease
- ( ) Don't know

9. 8.1) To your knowledge, has there been any change in hunting pressure faced by the following wildlife groups during the lock-down at this location? *

*Mark only one oval per row.*

| | Not hunted | Increased | Same | Decreased | Don't know |
|---|---|---|---|---|---|
| Mammals | ⬭ | ⬭ | ⬭ | ⬭ | ⬭ |
| Birds | ⬭ | ⬭ | ⬭ | ⬭ | ⬭ |
| Reptiles and amphibians | ⬭ | ⬭ | ⬭ | ⬭ | ⬭ |
| Fishes and crustaceans | ⬭ | ⬭ | ⬭ | ⬭ | ⬭ |

10. 8.2) Please share any details you have on what species were hunted, methods of hunting (snares, guns, etc) and why they were hunted (e.g. bushmeat species like deer and wild boar for local consumption etc.)

_______________________________________________

_______________________________________________

_______________________________________________

_______________________________________________

_______________________________________________

11. 9) Where did the hunting during lock-down take place? (Tick all applicable options) *

*Tick all that apply.*

- [ ] No hunting at this location
- [ ] Protected areas (National Parks, Wildlife Sanctuaries, etc.)
- [ ] Reserve forest (including community forest)
- [ ] Territorial forest
- [ ] Village revenue land
- [ ] Privately owned land
- [ ] Don't know

Other: [ ] _______________________

12. 10.1) How do you think the following motivations for hunting changed during the lock-down at this location? *

*Mark only one oval per row.*

| | Not a motivation | Increased | Same as before | Decreased | Don't know |
|---|---|---|---|---|---|
| Hunting for household consumption | ⬭ | ⬭ | ⬭ | ⬭ | ⬭ |
| Sale of meat in local markets | ⬭ | ⬭ | ⬭ | ⬭ | ⬭ |
| Medicinal use | ⬭ | ⬭ | ⬭ | ⬭ | ⬭ |
| Supply to outside markets | ⬭ | ⬭ | ⬭ | ⬭ | ⬭ |
| Sport and recreation | ⬭ | ⬭ | ⬭ | ⬭ | ⬭ |
| Trade in body parts | ⬭ | ⬭ | ⬭ | ⬭ | ⬭ |

13. 10.2) List any other motivations not listed above that you feel led to hunting at your location during the lock-down (e.g. hunting festival that coincided with the lock-down)

14. 11) Which groups, according to you, were responsible for hunting during the lock-down? (Tick all applicable options) *

*Tick all that apply.*

- [ ] No hunting at this location
- [ ] Resident who were regularly hunting at this location even before the lock-down
- [ ] Residents who were unemployed due to lock-down
- [ ] Individuals who moved back to this location during lock-down
- [ ] Outsiders and visitors to the location
- [ ] Mixed groups
- [ ] Don't know

Other: [ ] ______________________

15. 12) Select the local factors that may have contributed to an increase in hunting at your location? (Tick all applicable options) *

*Tick all that apply.*

- [ ] No increase in hunting at this location
- [ ] Lack of incomes from tourism, handicrafts, and other local industries
- [ ] Disruption of regular food supply
- [ ] Collapse of traditional seasonal occupations
- [ ] Lack of enforcement
- [ ] Need to supplement household income to sustain an influx of individuals from urban areas
- [ ] Need for recreational activity
- [ ] Community bonding (over hunting)
- [ ] Don't know

Other: [ ] ______________________

16. 13) How was enforcement against wildlife crime affected during the lock-down at this location? *

*Mark only one oval per row.*

| | Not present at this location | Increased | Same as before | Decreased | Don't know |
|---|---|---|---|---|---|
| Enforcement by Forest Department | ◯ | ◯ | ◯ | ◯ | ◯ |
| Enforcement by Police | ◯ | ◯ | ◯ | ◯ | ◯ |
| Community-patrolles (e.g. by village councils) | ◯ | ◯ | ◯ | ◯ | ◯ |
| Support from para-military forces | ◯ | ◯ | ◯ | ◯ | ◯ |
| Customs Agencies | ◯ | ◯ | ◯ | ◯ | ◯ |

17. 14) What do you think were the major challenges for enforcement during the lock-down? (Tick all applicable options) *

*Tick all that apply.*

- [ ] None
- [ ] Increased incidence of hunting
- [ ] Increased local support for hunting
- [ ] Lack of manpower for patrolling and other enforcement efforts
- [ ] Lack of mobility for enforcement staff
- [ ] Logistical challenges faced by the enforcement staff
- [ ] Difficulty in reaching information to enforcement agencies
- [ ] Enforcement staff were mobilized for other duties
- [ ] Don't know

Other: [ ] ____________________

18. 15.1) To your knowledge, which of these nongovernmental groups contribute towards controlling hunting at this location? (Tick all applicable options) *

*Tick all that apply.*

| | Did not contribute | Pre Lock-down | During Lock-down | Don't Know |
|---|---|---|---|---|
| Voluntary village groups | ☐ | ☐ | ☐ | ☐ |
| NGOs | ☐ | ☐ | ☐ | ☐ |
| Private companies (including tourism) | ☐ | ☐ | ☐ | ☐ |
| Private individuals | ☐ | ☐ | ☐ | ☐ |
| None | ☐ | ☐ | ☐ | ☐ |

19. 15.2) Please describe any other groups not listed above that are involved in controlling hunting at this location.

20. 16.1) What strategies were implemented by government and nongovernmental groups to control hunting at this location during the lock-down, and did they work? (Tick all applicable options) *

*Mark only one oval per row.*

| | Don't know if it was implemented | Not implemented | Implemented & Worked | Implemented & Did not work | Implemented & Don't know if it worked |
|---|---|---|---|---|---|
| More enforcement | ◯ | ◯ | ◯ | ◯ | ◯ |
| Provisioning of essential food supplies | ◯ | ◯ | ◯ | ◯ | ◯ |
| Monetary assistance to those who have lost jobs | ◯ | ◯ | ◯ | ◯ | ◯ |
| Awareness linking hunting and zoonotic diseases like COVID-19 | ◯ | ◯ | ◯ | ◯ | ◯ |
| Communication between enforcement agencies and local residents | ◯ | ◯ | ◯ | ◯ | ◯ |
| More logistic support to enforcement agencies. | ◯ | ◯ | ◯ | ◯ | ◯ |

21. 16.2) Please describe any other strategies not listed above that were implemented to reduce hunting at this location during the lock-down.

22. 17) Please list any additional observations you wish to share, including information on specific incidents (species involved, location, enforcement response, web-links to media coverage, etc.)

This content is neither created nor endorsed by Google.

Google Forms
