## Supplementary Material 2 for "Key informant perceptions on wildlife hunting in India during the COVID-19 lockdown"

***Table 1***. Overall changes in hunting

| **Change** | **Mean (%)** | **95% CI (Direct)** |
| --- | --- | --- |
| Don’t know | 27.23 | 19.19 – 36.36 |
| Increase | 56.12 | 40.33 – 74.54 |
| Decrease | 6.03 | 0.77 – 13.03 |
| No Change | 10.15 | 5.05 – 16.16 |

***Table 2.*** Change in hunting of taxa

| **Species** | **Change** | **Mean (%)** | **95% CI (Direct)** |
| --- | --- | --- | --- |
| Mammal | Same | 7.05 | 2.02 – 12.12 |
|  | Not Hunted | 15.13 | 8.08 – 22.22 |
|  | Increased | 55.10 | 45.46 – 65.66 |
|  | Decreased | 5.04 | 1.01 – 10.01 |
|  | Don’t know | 17.20 | 10.01 – 25.25 |
| Birds | Same | 16.12 | 9.09 – 23.23 |
|  | Not Hunted | 21.22 | 13.13 – 29.30 |
|  | Increased | 35.30 | 26.26 – 44.44 |
|  | Decreased | 5.04 | 1.01 – 10.01 |
|  | Don’t know | 22.28 | 14.14 – 31.31 |
| Rept. Amph. | Same | 18.19 | 11.11 – 26.26 |
|  | Not Hunted | 30.27 | 21.21 – 39.39 |
|  | Increased | 15.11 | 8.08 – 22.22 |
|  | Decreased | 2.02 | 0 – 5.05 |
|  | Don’t know | 34.41 | 25.25 – 43.43 |
| Fish Crust. | Same | 13.10 | 7.07 – 20.20 |
|  | Not Hunted | 12.12 | 6.06 – 19.19 |
|  | Increased | 43.40 | 34.34 – 53.54 |
|  | Decreased | 5.08 | 1.01 – 10.01 |
|  | Don’t know | 26.18 | 18.18 – 35.35 |

***Table 3***. Who is hunting

| **Who** | **Mean (%)** | **95% CI (Direct)** |
| --- | --- | --- |
| Resident Hunters | 63.62 | 54.54 – 72.72 |
| Resident (out of work) | 39.34 | 29.29 – 49.49 |
| Returnees | 20.24 | 12.12 – 28.28 |
| No hunting | 18.16 | 11.11 – 25.25 |
| Mixed Groups | 17.12 | 10.09 – 24.24 |
| Don’t know | 10.11 | 5.04 – 16.16 |
| Outsiders | 6.05 | 2.02 – 11.11 |

***Table 4.*** Where hunting took place

| **Location** | **Mean** | **95% CI (Direct)** |
| --- | --- | --- |
| Reserve Forest | 43.40 | 34.34 – 53.53 |
| Village Land | 32.32 | 23.23 – 41.41 |
| Protected Area | 28.26 | 19.19 – 37.37 |
| Private Land | 27.30 | 18.18 – 36.36 |
| Territorial Forest | 22.20 | 14.14 – 31.31 |
| No hunting | 20.23 | 12.12 – 28.28 |
| Don’t Know | 8.13 | 3.03 – 14.14 |

***Table 5.*** Changes in motivation to hunt

| **Motivation** | **Change** | **Mean (%)** | **95% CI (Direct)** |
| --- | --- | --- | --- |
| Household  Consumption | Same | 13.13 | 7.07 – 20.20 |
|  | Not a motive | 7.13 | 2.02 – 12.12 |
|  | Increased | 53.15 | 43.43 – 63.03 |
|  | Decreased | 5.06 | 1.01 – 10.10 |
|  | Don’t know | 21.24 | 13.13 – 29.29 |
| Local market | Same | 17.18 | 10.10 – 25.25 |
|  | Not a motive | 25.23 | 17.17 – 34.34 |
|  | Increased | 14.12 | 8.08 – 21.21 |
|  | Decreased | 14.16 | 8.08 – 21.21 |
|  | Don’t know | 29.30 | 20.20 – 38.38 |
| Medicine | Same | 14.12 | 8.08 – 21.21 |
|  | Not a motive | 31.35 | 22.22 – 40.40 |
|  | Increased | 12.12 | 6.06 – 19.19 |
|  | Decreased | 2.98 | 0 – 7.07 |
|  | Don’t know | 39.43 | 30.03 – 49.49 |
| Outside Market | Same | 9.13 | 4.04 – 15.15 |
|  | Not a motive | 30.25 | 21.21 – 39.39 |
|  | Increased | 11.05 | 5.05 – 17.17 |
|  | Decreased | 10.07 | 4.04 – 16.16 |
|  | Don’t know | 39.45 | 30.03 – 49.49 |
| Sports/Recreation | Same | 7.02 | 3.03 – 12.12 |
|  | Not a motive | 24.25 | 16.16 – 33.33 |
|  | Increased | 34.36 | 25.25 – 43.43 |
|  | Decreased | 6.08 | 2.02 – 11.11 |
|  | Don’t know | 28.26 | 19.19 – 37.37 |
| Trade | Same | 13.12 | 7.07 – 20.20 |
|  | Not a motive | 34.30 | 25.25 – 43.43 |
|  | Increased | 5.07 | 1.01 – 10.01 |
|  | Decreased | 4.04 | 1.01 – 8.08 |
|  | Don’t know | 43.49 | 34.32 – 53.53 |

***Table 6.*** *Local factors causing increase in hunting*

| **Changes** | **Mean (%)** | **95% CI (Direct)** |
| --- | --- | --- |
| Lack of enforcement | 36.40 | 27.27 – 45.45 |
| Food Disruption | 32.31 | 23.23 – 41.41 |
| Recreation | 32.31 | 23.23 – 41.41 |
| Seasonal Occupation Loss | 24.18 | 16.16 – 33.33 |
| No Tourist | 21.17 | 14.14 – 29.30 |
| No Increase | 19.19 | 12.12 – 27.27 |
| Community Bonding | 18.16 | 11.11 – 26.26 |
| Don’t Know | 12.14 | 6.06 – 19.19 |
| Supplementary Income | 7.05 | 2.02 – 12.12 |
| Others | 6.07 | 2.02 – 11.11 |

***Table 7***. Overall changes in enforcement

| **Enforcement** | **Change** | **Mean (%)** | **95% CI (Direct)** |
| --- | --- | --- | --- |
| Forest Department | Don’t know | 15.13 | 8.08 – 22.22 |
|  | Not present | 18.20 | 11.11 – 26.26 |
|  | Same as before | 34.34 | 25.25 – 44.44 |
|  | Decreased | 12.15 | 6.06 – 19.19 |
|  | Increased | 20.19 | 12.12 – 28.28 |
| Police Department | Don’t know | 26.29 | 18.18 – 35.35 |
|  | Not present | 28.33 | 19.19 – 37.37 |
|  | Same as before | 29.26 | 20.20 – 38.38 |
|  | Decreased | 7.11 | 3.03 – 12.12 |
|  | Increased | 9.13 | 4.04 – 15.15 |
| Customs | Don’t know | 39.27 | 30.03 – 48.48 |
|  | Not present | 45.40 | 35.35 – 55.56 |
|  | Same as before | 9.09 | 4.04 – 15.15 |
|  | Decreased | 5.08 | 1.01 – 10.01 |
|  | Increased | 1.02 | 0 – 3.03 |
| Community | Don’t know | 31.26 | 22.22 – 40.04 |
|  | Not present | 39.39 | 29.29 – 49.49 |
|  | Same as before | 12.10 | 6.06 – 19.19 |
|  | Decreased | 10.12 | 5.05 – 16.16 |
|  | Increased | 7.07 | 2.02 – 12.12 |
| Paramilitary | Don’t know | 38.43 | 29.29 – 48.48 |
|  | Not present | 49.48 | 39.39 – 59.60 |
|  | Same as before | 7.11 | 2.02 – 12.12 |
|  | Decreased | 3.01 | 0 – 7.07 |
|  | Increased | 1.98 | 0 – 5.05 |

***Table 8***. Challenges to enforcement

| **Changes** | **Mean (%)** | **95% CI (Direct)** |
| --- | --- | --- |
| Lack of Manpower | 49.59 | 39.39 – 59.60 |
| Logistical Challenges | 38.44 | 29.29 – 48.48 |
| Lack of mobility | 38.44 | 29.29 – 48.48 |
| Increased Hunting | 36.37 | 27.27 – 45.45 |
| Communication | 27.27 | 19.19 – 36.36 |
| Increased local support | 27.27 | 19.18 – 36.35 |
| Don’t Know | 14.11 | 6.67 – 19.10 |
| None | 13.22 | 7.07 – 20.20 |
| Diversion | 13.22 | 7.07 – 20.20 |
| Others | 6.04 | 1.01 – 11.10 |

***Table 9***. Strategies to deal with hunting

| **Strategy** | **Change** | **Mean (%)** | **95% CI (Direct)** |
| --- | --- | --- | --- |
| More Enforcement | Don’t know | 37.37 | 28.28 – 47.47 |
|  | Implemented but didn’t work | 4.05 | 1.01 – 8.08 |
|  | Implemented but don’t know if it worked | 15.14 | 8.08 – 22.22 |
|  | Implemented & worked | 13.12 | 7.07 – 20.20 |
|  | Not implemented | 30.20 | 21.21 – 39.39 |
| Provision Food | Don’t know | 34.35 | 25.25 – 43.43 |
|  | Implemented but didn’t work | 8.08 | 3.03 – 14.14 |
|  | Implemented but don’t know if it worked | 22.24 | 14.14 – 30.30 |
|  | Implemented & worked | 17.24 | 10.10 – 25.25 |
|  | Not implemented | 18.17 | 11.11 – 26.26 |
| Monetary Assistance | Don’t know | 46.39 | 36.36 – 56.57 |
|  | Implemented but didn’t work | 5.02 | 1.01 – 10.10 |
|  | Implemented but don’t know if it worked | 10.11 | 5.05 – 16.16 |
|  | Implemented & worked | 4.01 | 1.01 – 8.08 |
|  | Not implemented | 34.32 | 25.25 – 43.43 |
| Awareness | Don’t know | 35.43 | 26.26 – 44.44 |
|  | Implemented but didn’t work | 5.05 | 1.01 – 10.01 |
|  | Implemented but don’t know if it worked | 11.11 | 5.05 - 17.17 |
|  | Implemented & worked | 8.02 | 3.03 – 14.14 |
|  | Not implemented | 40.29 | 30.30 – 49.50 |
| Communication | Don’t know | 36.35 | 27.27 – 45.45 |
|  | Implemented but didn’t work | 9.05 | 4.04 – 15.15 |
|  | Implemented but don’t know if it worked | 15.17 | 8.08 – 22.22 |
|  | Implemented & worked | 8.12 | 3.03 – 14.14 |
|  | Not implemented | 31.37 | 22.22 – 40.04 |
| Logistic Support | Don’t know | 46.51 | 37.37 – 56.57 |
|  | Implemented but didn’t work | 5.06 | 1.01 – 10.01 |
|  | Implemented but don’t know if it worked | 11.11 | 5.05 – 17.17 |
|  | Implemented & worked | 6.08 | 2.02 – 11.11 |
|  | Not implemented | 31.25 | 22.22 – 40.40 |

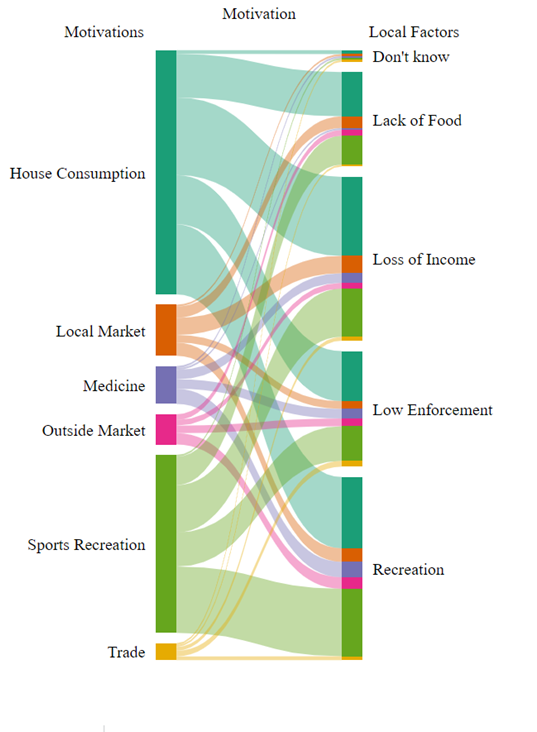

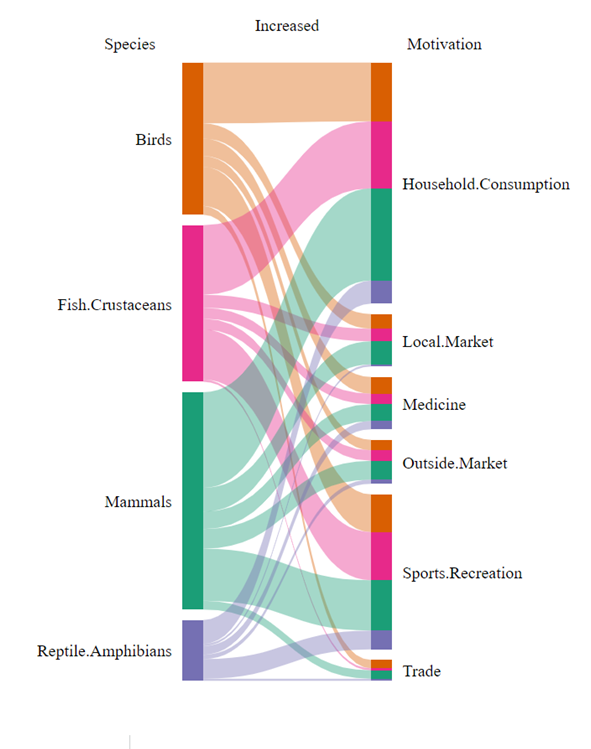

***Fig 1.*** *Left: a. Association between the change in motivations to hunt and the lockdown-specific local factors were perceived to have resulted in an increase*. Local factors were binned from the set displayed in figure 4C. *Right: b*. *Association between the change in motivations to hunt and change in hunting pressure on different taxa during lockdown*
