## Supplementary Material 3.2 for "Key informant perceptions on wildlife hunting in India during the COVID-19 lockdown"

**Media Analysis Methods and Results**

Important note on methodology:

1. Three set of search phrases: “Lockdown, hunting, wildlife, killing”; “COVID-19, lockdown, hunting” ; “COVID-19, wildlife, hunting, poaching, India” were used to search for media articles on search engines
2. Any inventory of all the articles were made. Duplicates were removed. (see Media article sheet in the excel)
3. For each article, statements(direct or indirect quotes) by experts/key informants on hunting during the lockdown were highlighted (see Media article sheet in the excel)
4. For each quote: who said the quote, motivation behind hunting, information regarding local factors, enforcement, challenges to enforcement, location, species and overall trend in hunting was noted (if available; see Qoutes analyzed sheet in the excel)
5. Same quotes by the same key- informant/expert was only considered once to ensure no pseudo replication.

Below are a summary of results of the media analysis

Total Number of articles obtained: **119**

Total unique articles analyzed (Criteria = before 31^st^ May 2020): **98**

**Hunting Trends**

| **Trends** | **No. of Statement** | **No. of unique experts** | **Animals** | | | | | | **States** |
| --- | --- | --- | --- | --- | --- | --- | --- | --- | --- |
|  |  |  | **Ungulate** | **Lagomorph** | **Rodent** | **Other mammals** | **Birds** | **Reptiles** |  |
| Increase | 78 | 60 | Barking Deer, Chinkara, Blackbuck, Nilgai, Spotted Deer, Wild Board, Sambar, One-horned Rhino, Gaur  (9) | Desert Hare, Black-Naped Hare  (2) | Porcupine , Giant Squirrel sp.  (2) | Asian Palm Civet, Large India Civet, Pangolin sp., Leopard  (4) | Grey Francolin, Peacock, Jungle Fowl, H. Monal, Moorhen, Parrot sp.  (6) | Python, King Cobra, Monitor Lizard, Spiny-tailed Lizard, Crocodile  (5) | Arunachal Pradesh, Assam, Himachal Pradesh, Karnataka, Kerala, Madhya Pradesh, Maharashtra, Nagaland, Odisha, Rajasthan, Tamil Nadu, Uttarakhand |
| Decrease | 3 | 3 | - | - | - | Tiger | - | - | Goa, Nagaland |
| Same | 10 | 10 | Wild Boar, Spotted Deer, Sambar, Gaur |  |  | Tiger, Pangolin |  | King Cobra | Andhra Pradesh, Arunachal Pradesh, Goa, Karnataka, Kerala, Maharashtra |
| No hunting | 4 | 2 | - | - | - | - | - | - | Andrha Pradesh, Tamil Nadu |

**Table 1.** Table displays the various statements with respect to the trends of hunting during lockdown. Each trend is linked to its corresponding animals sp. and the state from which this information is obtained.

**Motivation of hunting – Increase**

**Table 2.** Table displays the various motivations and the local factors driving those motivations that lead to increase in hunting for certain species in certain states during the lockdown.

| **Motivation *‡** | **Local factors**  **(statements)** | **Species** | **State** |
| --- | --- | --- | --- |
| **Household Consumption**  * =18  **‡ =**16 | Disruption of regular food supply (9) | King Cobra | Arunachal Pradesh |
|  |  | Chinkara | Rajasthan |
|  |  | Spotted Deer | Karnataka |
|  |  | Nilgai, Spotted Deer | Madhya Pradesh |
|  |  | Grey Francolin | Rajasthan |
|  |  | Tiger’s prey | - |
|  | Lack of Income (5) | Nilgai, Spotted Deer, Wild Boar | Maharashtra |
|  |  | Chinkara | Rajasthan |
|  |  | Spotted Deer, Pangolin, Wild Boar | Odisha |
|  |  | Himalayan Monal | Himachal Pradesh |
|  | Influx of people form urban areas (1) | Spotted Deer, Pangolin, Wild Boar | Odisha |
|  | Recreational needs (5) | Himalayan Monal | Himachal Pradesh |
|  |  | Barking Deer and Large Indian Civet | Nagaland |
|  | Don’t know (4) | Moorhen and Parrots | Kerala |
|  |  | Moorhen | Assam |
|  |  | Hare sp. | Tamil Nadu |
| **Sports & Recreation**  * =6  **‡ =**5 | Recreational needs (5) | Barking Deer, Large Indian Civet, Python | Nagaland |
|  |  | Deer, Wild Boar, Monitor Lizard, Jungle Fowl, Porcupine, Black Naped Hare | Tamil Nadu |
|  | Community Bonding (1) | Barking Deer, Python | Nagaland |
|  | Don’t Know (1) | - | Nagaland |
| **Sale of meat in local markets**  * =3  **‡ =**3 | Disruption of regular food supply (1) | Spotted Deer, Sambar, Wild Boar | - |
|  | Lack of income (2) | Spotted Deer, Sambar, Wild Boar |  |
|  |  | Spotted Deer, Pangolin, Wild Boar | Odisha |
|  | Influx of people from urban areas (1) | Spotted Deer, Pangolin, Wild Boar | Odisha |
|  | Don’t know (1) | Moorhen | Assam |
| **Trading in body Parts**  * = 2 ; **‡ =** 2 | Post Lockdown demand (1) | One-horned Rhino | Assam |
|  | Don’t know (1) | One-horned Rhino | Assam |
| **Don’t know**  * =52  **‡ =**45 | Disruption of regular food supply (3) | Crocodile, Blackbuck, Nilgai, Spotted Deer, Chinkara, Wild boar, Wild Hare, Peacock | Madhya Pradesh |
|  |  | - | Karnataka |
|  |  | Leopard | Nagaland |
|  | Lack of income (3) | Leopard | Nagaland |
|  |  | - | Rajasthan |
|  | Influx of people from urban areas (3) | - | Rajasthan |
|  |  | Pangolin and Porcupine | Uttarakhand |
|  | Recreational needs (3) | Moorhen | Assam |
|  |  | - | Arunachal Pradesh |
|  |  | - | Karnataka |
|  | Lack of enforcement (1) | Peacock | Madhya Pradesh |
|  | Reduced local movement (2) | Wild boar, rabbits, deer, peacock | Karnataka |
|  |  | Chinkara, Blackbuck | Rajasthan |
|  | Don’t know (38) | *Arunachal Pradesh* = NA; *Assam* = One-horned Rhino, Moorhen; *Karnataka* = Spotted Deer, Sambar, Wild Boar, Peacock; *Madhya Pradesh* = Nilgai; *Maharashtra* = Peacock, Rabbits, Wild Boar, Barking Deer, Civets, Leopard, Chinkara; *Nagaland* =NA, *Odisha* = Pangolins; Rajasthan = Chinkara, Blackbuck, Spiny-tailed Lizard, Desert Hare, Peafowl, Monitor Lizard, Grey Francolin, Nilgai, *Tamil Nadu* = Jungle Fowl, Wild boar, Black-naped hare, Sambar and Barking deer, Gaur, leopard, Tiger, Monitor Lizard. | |

* = Number of unique statements; **‡ =** Number of unique experts

Note: The colors are kept similar for the same local factors driving various motivations. For instance red is for “Disruption of regular food supply”.

**Enforcement – Overall**

**Table 3.** How the enforcement levels of authorities changes during lockdown, major challenges they face and the trend in hunting that given state. Numbers in the brackets are the unique statements.

| **Enforcement authority** | **Enforcement level** | **Major Challenges** | **Trend in Hunting** | **States** |
| --- | --- | --- | --- | --- |
| **Community Patrols**  (15) | Same (6) | Lack of manpower(1) | Same (1) | Goa |
|  |  | - | Decreased (2) | Goa, Nagaland |
|  |  |  | Don’t know (3) | Uttarakhand, Maharashtra, Assam |
|  | Increased (6) | Lack of manpower (1) | Don’t know (1) | Goa |
|  |  | - | Increase (2) | Rajasthan |
|  |  | - | Don’t know (3) | Goa, Rajasthan |
|  |  | - | No hunting (1) | Andhra Pradesh |
|  | Decreased (3) | Lack of mobility (2) | Increase (1), Don’t know (1) | Maharashtra, Odisha |
|  |  | Enforcement has other duties (1) | Increase (1) | - |
| **Forest Department**  (34) | Same (10) | Enforcement has other duties (1) | - | Maharashtra |
|  |  | - | Increase (1) | Karnataka |
|  |  | - | Don’t know (9) | Arunachal Pradesh, Uttarakhand, Himachal Pradesh, Maharashtra, Kerala, Tamil Nadu, Karnataka |
|  | Increased (18) | None (1) | Don’t know(1) | Rajasthan |
|  |  | - (16) | Increase (2) | Arunachal Pradesh, Tamil Nadu |
|  |  |  | Don’t know (14) | UP, Uttarakhand, HP, Karnataka, Tamil Nadu, Odisha, Rajasthan, Kerala |
|  |  | - (1) | No hunting (1) | Andhra Pradesh |
|  | Decreased (7) | Lack of mobility (3) | Don’t know (2) | Assam, UP, Uttarakhand |
|  |  |  | Increase (1) | Odisha |
|  |  | Logistical challenges (1) | Increase (1) | General |
|  |  | Enforcement other duties (2) | Increase (2) | General, Assam |
|  |  | - (1) | Increase (1) | Nagaland |
| **Police Department**  (3) | Increase (2) | - (2) | Don’t know (2) | HP, Kerala |
|  | Decrease(1) | Enforcement other duties (1) | Increase (1) | - |
